# Modifications of Two ESCRT-I Subunits with Distinct Ubiquitin Chains Regulate Plant Immunity

**DOI:** 10.1101/2023.08.10.552820

**Authors:** Chaofeng Wang, Yi Zhang, Bangjun Zhou, Pooja Verma, Sadia Hamera, Lirong Zeng

## Abstract

Sensing of pathogen- or microbe-associated molecular patterns (PAMPs/MAMPs) by pattern recognition receptors (PRRs) at the cell surface induces the first layer of host immunity against invading microbial pathogens. The immune receptor FLS2 perceives bacterial flagellin to initiate host immune signaling upon pathogen infections. It has been known that the FLS2 abundance is crucial for plant pattern-triggered immunity. Nevertheless, the underpinning regulatory mechanisms remain largely unclear. In this study, we demonstrate that XBAT35.2 positively modulates the protein level of FLS2. In addition to the Golgi, XBAT35.2 localizes at the plasma membrane and constitutively associates with FLS2, BAK1, and BIK1. Flg22 treatment increases the association of XBAT35.2 with FLS2 and BAK1 but reduces the interaction with BIK1. XBAT35.2 ubiquitinates two key components of the ESCRT-I complex, VPS37-1 and VPS28-2 with K48 and K63-linked polyubiquitin chains respectively, leading to degradation of VPS37-1 and diminished interaction of VPS28-2 with FLS2. Additionally, VPS37-1 and VPS28-2 play redundant, negative roles in FLS2-mediated immunity by promoting vacuolar breakdown of FLS2. Thus, by intercepting the function of VPS28-2 and VPS37-1, XBAT35.2 stabilizes FLS2 for host immunity. Our findings reveal a new regulatory circuit for modulating the FLS2 abundance and deepen our understanding of controlling the homeostasis of cell surface receptors.

**One-sentence summary:** Targeting the ESCRT-I subunits VPS28-2 and VPS37-1 with distinct ubiquitin chains by the E3 ligase XBT35.2 stabilizes FLS2 and positively modulates plant immunity.

## Introduction

Plants and animals employ innate immunity to prevent infections by recognition of PAMPs/MAMPs using cell surface-resident pattern-recognition receptors (PRRs) (Constant et al., 2021; DeFalco and Zipfel, 2021; Ngou et al., 2022). Detection of PAMPs/MAMPs activates pattern-triggered immunity (PTI), the first layer of a multitier host innate immune system (Ngou et al., 2022). Plant PRR Flagellin Sensing 2 (FLS2) is a plasma membrane (PM)-localized leucine-rich repeat receptor-like kinase (LRR-RLK) that recognizes bacterial flagellin and flg22, an immunogenic 22-amino acid peptide derivative of flagellin. Perception of flagellin/flg22 activates FLS2 and instantaneously leads to the association of FLS2 with Brassinosteroid-Insensitive 1- Associated Kinase 1 (BAK1), another PM-localized LRR-RLK that serves as co-receptor for multiple PRRs. This affiliation of FLS2 with BAK1 simultaneously results in the formation of a heterotrimer complex that includes FLS2, BAK1, and Botrytis-Induced Kinase 1 (BIK1), a PM-associated receptor-like cytoplasmic kinase (RLCK). Early signaling events in the FLS2-BAK1-BIK1 complex include sequential autophosphorylation and transphosphorylation within the heterotrimer and mono-ubiquitination of BIK1, which leads to the release of BIK1 from the complex and sophisticated downstream signaling cascades that culminate in various immune responses (Zhou and Zhang, 2020; DeFalco and Zipfel, 2021; Kong et al., 2021).

FLS2-mediated PTI plays a critical role in plant defense against bacterial pathogens, including the *Pseudmonas syringae* pv *tomato* (*Pst*) strain DC3000 (Zipfel et al., 2004; Rosli et al., 2013). Despite the importance, however, excessive or ill-timed activation of innate immune responses is detrimental to the host (Dominguez-Ferreras et al., 2015; Cao, 2016). Conceivably, the abundance and activity of PRRs must be fine-tuned and tightly controlled such that the amplitude and duration of PTI responses are precisely adjusted. The abundance of FLS2 has hitherto been shown to be regulated by both ubiquitin-26S proteasome-dependent protein turnover and endocytosis-mediated recycling/degradation. The former involves ubiquitination of FLS2 by two closely related *Arabidopsis* U-box type ubiquitin E3 ligases, PUB12 and PUB13 (Lu et al., 2011). In the presence of flagellin/flg22, BAK1 phosphorylates PUB12 and PUB13. Phosphorylated PUB12 and PUB13 physically interact with and ubiquitinate FLS2 to promote degradation by the 26S proteasome, which attenuates FLS2-mediated immune signaling (Lu et al., 2011). Endocytosis and the following intracellular endosomal trafficking occur on both nonactivated and flagellin/flg22-activated FLS2, targeting the receptor for recycling to PM and for degradation probably at the vacuoles, respectively (Robatzek et al., 2006; Beck et al., 2012). Endocytic recycling of nonactivated FLS2 takes place constitutively between the PM and the *trans*-Golgi network (TGN)/early endosome (EE) compartments and is believed to maintain a constant pool of FLS2 at the PM (Beck et al., 2012). By contrast, activated FLS2 proteins are internalized and then routed through TGN/EEs, intermediate compartments with characteristics between the TGN/EE and late endosome (LE) /multivesicular body (MVB), and LE/MVB compartments or pre-vacuolar compartments (PVCs) before putative vacuolar breakdown (Robatzek et al., 2006; Lu et al., 2011; Beck et al., 2012; Choi et al., 2013; Paez Valencia et al., 2016). The predominant route of constitutive and ligand-induced internalization of cell surface receptors in plants is mediated by clathrin-dependent endocytosis, a complex process executed through spatiotemporal coordination of multiple factors, such as dynamin GTPase proteins (Smith et al., 2014; Paez Valencia et al., 2016; Ekanayake et al., 2021). Consistently, the internalization of ligand-activated FLS2 was shown to be dependent on the clathrin protein that is essential for immunity against bacterial infection (Mbengue et al., 2016). The *Arabidopsis* dynamin-related protein 2B (DRP2B) was found to function in plant immune responses to flg22 and *Pst* DC3000 treatment (Smith et al., 2014). In the *drp2b* null mutant, flg22-induced endocytosis of FLS2 was partially reduced. More recently, it was revealed that DRP1A, a member of the DRP1 subfamily representing plant-specific dynamins, is also required for effective immunity against *Pst* DC3000 infection and for flg22-induced immune signaling (Ekanayake et al., 2021). DRP1A and DRP2B play synergistic roles in governing the abundance of FLS2 at PM by regulating its endocytosis. In addition to these components involved in the early stage of endocytosis, two subunits of the endosomal sorting complex required for transport (ESCRT)-I complex, vacuolar protein sorting (VPS) 28-2 and VPS37-1 were shown to be involved in the sorting and delivery of activated FLS2 to the MVB lumina (Spallek et al., 2013). During the endocytosis, a large portion of flg22-activated FLS2s were found to be in constant association with LE/MVB, the compartments where cargo proteins are frequently targeted for vacuolar breakdown (Beck et al., 2012; Choi et al., 2013). It is thus believed that endocytosis and the following endosomal trafficking/sorting of activated FLS2 also contribute to the attenuation of FLS2-mediated immunity. Other than these findings, the molecular mechanism underlying induction of the endocytosis of activated FLS2 and regulation of the ensuing endosomal trafficking and sorting remain poorly understood. In fact, despite their overarching roles in numerous pathways (Paez Valencia et al., 2016; Vietri et al., 2020; Sigismund et al., 2021), knowledge regarding how the machinery responsible for endocytosis and endosomal trafficking is modulated remains very limited.

Ubiquitination is one of the most abundant types of post-translational protein modifications in eukaryotes and plays key roles in modulating plant immunity (Zhou and Zeng, 2017; Zhang and Zeng, 2020). Through a stepwise enzymatic cascade that involves three different classes of enzymes, ubiquitin-activating enzyme (E1 or UBA), ubiquitin-conjugating enzyme (E2 or UBC), and ubiquitin ligase (E3), ubiquitination covalently attaches ubiquitin molecule to a lysine (K) residue of the substrate protein. After the first ubiquitin moiety is attached to the substrate protein, free ubiquitin molecules can be continuously linked to the precedent ubiquitin to assemble various types of poly-ubiquitin chains through one of the seven lysine residues (K6, K11, K27, K29, K33, K48, and K63) within the ubiquitin molecule. Canonically, ubiquitination is referred to as formation of K48-linked polyubiquitin chain, which is the most abundant type of poly-ubiquitination and is recognized as a principal signal for degradation of targeted proteins via the 26S proteasome. Nonetheless, the types of ubiquitination that occur in a cell are highly diverse and the importance of noncanonical ubiquitination in roles other than proteasome-dependent protein turnover have been increasingly appreciated (Zhou and Zeng, 2017). For instance, mono- or K63-linked ubiquitination serves as a key signal for guiding internalization and ensuing trafficking of PM-resident proteins to the lysosome/vacuole for degradation (Paez Valencia et al., 2016). It has long been shown that the ubiquitin system is essential for endocytosis of activated FLS2 (Robatzek et al., 2006). However, it remains unclear whether the involvement of ubiquitination in FLS2 endocytosis happens at the initial internalization step or the ensuing cargo trafficking and sorting steps, or both. Neither is known if noncanonical ubiquitination, especially K63-linked ubiquitination plays a role in modulating the trafficking and sorting machinery that is critical for endocytosis of various cell-surface receptors.

In this study, we demonstrate that the Ankyrin repeats (ANK)-RING type ubiquitin E3 ligase XBAT35.2 positively regulates FLS2-mediated plant innate immunity against the bacterial pathogen *Pst* DC3000. Our results indicate that XBAT35.2 modulates the protein level of FLS2 and specifically interacts with and catalyze attachment of K48 and K63-linked polyubiquitin chains to the two key ESCRT-I subunits, VPS37-1 and VPS28-2, respectively, which leads to proteasomal degradation of VPS37-1 and reduced interaction of VPS28-2 with FLS2. Through sabotaging VPS28-2/VPS37-1-mediated vacuolar degradation of FLS2, XBAT35.2 stabilizes FLS2 to effectively mount immunity against bacterial pathogens. Given the overarching role of endocytosis and ensuing endosomal trafficking and sorting in controlling the protein level of cell surface receptors, our findings support the notion that modification of ESCRT machinery components by ubiquitin could be a widespread regulatory mechanism for cell surface receptors-mediated cellular signaling.

## Results

### Identification of <u>F</u>ni<u>3</u>-interacting (Fti) proteins in tomato

Recognition of the *Pseudomonas syringae* effector AvrPtoB variants by the protein kinase Fen plays an important role in tomato (*Solanum lycopersicum*) defense against bacterial infections (Rosebrock et al., 2007). We previously identified Fni3 (<u>F</u>e<u>n</u>-interacting protein <u>3</u>), a tomato UBC13-type E2 enzyme and revealed that Fni3 acted with its cofactor Suv to modulate Fen-mediated immune responses by K63–linked ubiquitination (Mural et al., 2013). To further our understanding of the roles for ubiquitination in regulating plant immunity, we performed yeast two-hybrid (Y2H) screens for <u>F</u>ni<u>3</u>-interacting (Fti) proteins against a tomato cDNA library prepared from leaves of tomato plants challenged with *Pst* DC3000 (Zhou et al., 1995). Eight Fti proteins were identified from these screens, among which over 40% of the positive clones harbor sequences encoding part or full length of Fti1 (<u>F</u>ni<u>3</u>-interacting protein <u>1</u>). BLAST search of the tomato genome revealed that Fti1 belongs to a small family of five members (we temporarily named them Fti1, Fti1B, Fti1C, Fti1D, and Fti1E, respectively) and Fti1 displays the highest homology to the Fti1B protein encoded by the gene at locus Solyc06g008500 (Supplementary Fig. 1a). Y2H assay indicated that, among the 5 family members, only Fti1 and Fti1B interact with Fni3 (Fig. 1a) and the interactions were confirmed by bimolecular fluorescence complementation (BiFC) assay in tomato protoplasts (Supplementary Fig. 1b).

**Figure 1.**
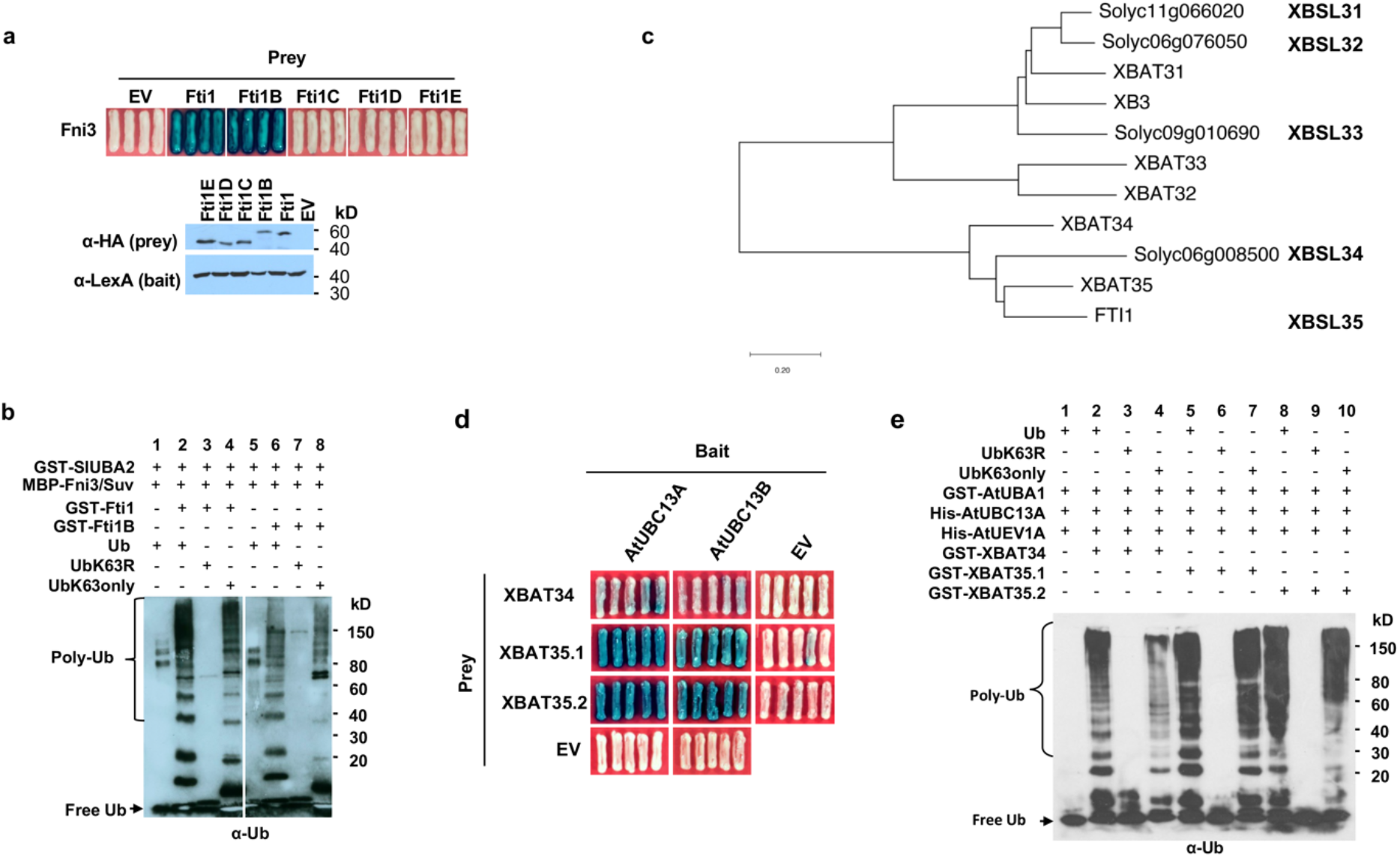
Tomato Fti1 (XBSL35), Fti1B (XBSL34) and their closest *Arabidopsis* homologs, XBAT35 and XBAT34 are active ubiquitin E3 ligases. **(a)** The interaction of Fni3 with Fti1 and other family members in yeast two-hybrid (Y2H) assay. Blue patches show positive interactions. Upper panel: photos were taken 24 h after yeast cells were grown on selection media supplied with X-gal. Lower panel: immunoblot shows expression of the bait (α-LexA) and prey (α-HA) proteins in the yeast cells. The numbers on the right show the molecular mass of marker proteins in kilodaltons. This experiment was repeated twice with similar results. **(b)** Fti1 and Fti1B work with the UBC13 type E2 Fni3 and its co-factor, Suv to direct K63-linked ubiquitination. The numbers on the right show the molecular mass of marker proteins in kilodaltons. This experiment was repeated three times with similar results. **(c)** Phylogenetic tree of tomato (*Solanum lycopersicum*) Fti proteins and their *Arabidopsis* homologs. Rice ANK-RING type protein XB3 was included in the phylogenetic analysis. The tree was constructed using the MEGA11 software (Tamura et al., 2021) by the neighbor-joining method with 500 bootstrap replicates. **(d)** XBAT34 and XBAT35 isoforms XBAT35.1 and XBAT35.2 interact with *Arabidopsis* UBC13A and UBC13B in Y2H. The blue patch denotes interaction of bait and prey proteins. Photos were taken 24 h after yeast cells were grown on selection media supplied with X-gal. This experiment was repeated twice with similar result. **(e)** *Arabidopsis* XBAT34 and XBAT35 work with UBC13A and its co-factor UEV1A to direct K63-linked ubiquitination. The numbers on the right show the molecular mass of marker proteins in kilodaltons. This experiment was repeated twice with similar results.

Further sequence analysis of Fti1 indicated it possesses a putative myristoylation site at the N-end, a N-terminal ankyrin repeats domain, a central putative ubiquitin interacting motif, and a C-terminal RING domain (Supplementary Fig. 2a and 2b). The ANK-RING domain organization of Fti1 and Fti1B is reminiscent of the rice XB3 and *Arabidopsis* XBAT3 proteins (Nodzon et al., 2004; Wang et al., 2006). Accordingly, we renamed the tomato Fti1 and Fti1B proteins as XBSL35 and XBSL34 respectively (for <u>XB</u>3 ortholog 5 and 4 in *<u>S</u>olanum lycopersicum*) based on the result that they show the highest homology to *Arabidopsis* XBAT35 and XBAT34, correspondingly (Fig. 1c).

### XBSL34, XBSL35 and their closest *Arabidopsis* homologs, XBAT34 and XBAT35 are active ubiquitin E3 ligase

Both XBSL35 and XBSL34 contain a C-terminal RING domain, suggesting that they might be active ubiquitin E3 ligases. Indeed, *in vitro* ubiquitination assays indicated that both XBSL35 and XBSL34 can catalyze autoubiquitination, as observed by the formation of high molecular mass proteins detected by immunoblotting analysis using anti-ubiquitin antibodies (Fig. 1b, lanes 2 and 6). As negative control, no polyubiquitination signals were detected when XBSL35 or XBSL34 were absent (Fig. 1b, lanes 1 and 5). The UBC13 type E2 Fni3 and Suv together were shown to assemble exclusively K63–specific polyubiquitination chains (Mural et al., 2013). We thus also examined the linkage types of the polyubiquitin chains formed in the autoubiquitination using different ubiquitin mutants in the *in vitro* ubiquitination assay. When the K63 residue was substituted with Arg (UbK63R), the formation of polyubiquitin chains was abolished (Fig. 1b, lanes 3 and 7). However, when the ubiquitin variant in which all K residues were substituted with R except only K63 remained intact (UbK63only) was used, XBSL35 and XBSL34 directed formation of polyubiquitin chains comparable to that of the wild-type ubiquitin (Fig. 1b, lanes 4 and 8). These findings indicate that XBSL35 and XBSL34 are *bona fide* ubiquitin E3 ligase and can work with Fni3/Suv to direct K63-linked ubiquitination.

In *Arabidopsis*, the *XBAT35* mRNA undergoes alternative splicing, producing two isoforms: *XBAT35.1* and *XBAT35.2* (Carvalho et al., 2012). The isoform XBAT35.1 contains a nuclear localization signal (NLS) sequence that is absent in XBAT35.2. Interestingly, we also obtained two transcript products when gene-specific primers were used for amplifying the tomato *XBSL35* gene (Supplementary Fig. 3a). Sequence analysis indicated that the longer product contains an NLS sequence that is absent in the shorter one, suggesting that the alternative splicing is conserved in the *Arabidopsis XBAT35* and tomato *XBSL35* gene. We thus named the longer product *XBSL35.1*, and the shorter one *XBSL35.2* (Supplementary Fig. 3b and 3c).

The high homology of the *Arabidopsis* XBAT34 and XBAT35 to the tomato XBSL34 and XBSL35 encouraged us to test whether the *Arabidopsis* XBAT34 and XBAT35 are also able to act with *Arabidopsis* UBC13-type E2s to direct assembly of K63-linked polyubiquitin chains. The *Arabidopsis* genome encodes two UBC13 type E2 proteins, UBC13A (also named AtUBC35) and UBC13B (AtUBC36) that are 98% identical in protein sequence. We first carried out Y2H assays and found that the two XBAT35 isoforms, XBAT35.1 and XBAT35.2 strongly interacted with UBC13A and UBC13B (Fig. 1d). The interactions between XBAT34 and UBC13A and UBC13B were also detected, albeit much weaker (Fig. 1d). We next performed *in vitro* ubiquitination assays like what we did with XBSL35 and XBSL34 (Fig. 1b). In the presence of E1 (AtUBA1), E2 (UBC13A), UEV1A, and intact ubiquitin, XBAT34, XBAT35.1 and XBAT35.2 could catalyze autoubiquitination (Fig. 1e, lanes 2, 5, and 8). When the intact ubiquitin was replaced by UbK63R, no polyubiquitin chain signals were detected (Fig. 1e, lanes 3, 6, and 9); however, when UbK63only was used, comparable ubiquitin-positive smears were also detected (Fig. 1e, lanes 4, 7, and 10), indicating that XBAT34 and XBAT35, like their tomato homolog XBSL34 and XBSL35, can work with UBC13 type E2 to direct K63-linked ubiquitination. We also showed that XBAT35.2 can work with the E2 UBC8 that usually directs K48-linked ubiquitination to catalyze autoubiquitination (Supplementary Fig. 2c), corroborating a previous report that XBAT35 directs assembly of K48-linked polyubiquitination chains (Yu et al., 2020). Thus, XBAT34/35, like XBSL34/35, are active E3 ligases that can work with UBC13 type E2 to direct K63-specific polyubiquitination, and XBAT35 also mediates K48-linked polyubiquitination.

### XBAT35.2 but not XBAT34 and XBAT35.1 functions in plant immunity against bacterial infection

The high homology between the tomato XBSL34, XBSL35, and the *Arabidopsis* XBAT34, XBAT35, the results that all the four proteins can work with UBC13 type E2 to catalyze K63-linked ubiquitination, and the conservation between *XBAT35* and *XBSL35* in mRNA alternative splicing prompted us to use XBAT34 and XBAT35 from *Arabidopsis* as a starting point for functional characterization of these ANK-RING type E3 ligases in plant immunity.

We first obtained T-DNA insertion mutants for *XBAT34* (AT4G14365), and *XBAT35* (AT3G23280), respectively referred to as *xbat34* (SALK_132378) and *xbat35-1* (SALK_104813) (Supplementary Fig. 4a). Considering the possible functional redundancy between XBAT34 and XBAT35, *xbat34/xbat35-1* double mutant lines were generated by crossing. Examination of *XBAT34* and *XBAT35* expression in these mutants indicated that the corresponding gene was disrupted (Supplementary Fig. 4b). No developmental or morphological defects under normal growth conditions were observed in these single and double mutants (Supplementary Fig. 4c). We then inoculated *xbat34*, *xbat35-1*, and *xbat34/35-1* plants with the bacterial pathogen *Pst* DC3000 and examined the pathogen growth 3 days post inoculation (dpi). *Pst* DC3000 growth on *xbat34* was comparable to that on Col-0, however, the bacterial growth on *xbat35-1* was significantly higher than that on Col-0 and *xbat34* (Fig. 2a). The double mutant *xbat34/35-1* showed a similar level of *Pst* DC3000 growth to that on the *xbat35-1* single mutant (Fig. 2a). These results indicated that XBAT35 plays a positive role but XBAT34 seemingly does not function in host immunity against *Pst* DC3000. To further confirm this, we obtained another allele of *xbat34* T-DNA insertion line, termed here *xbat34*-#57 (CS873757) (Supplementary Fig. 5a and 5b). The growth of *Pst* DC3000 at 2 dpi and 3 dpi on this mutant line also was comparable to that on Col-0 plants (Supplementary Fig. 5c). We thus focused on XBAT35 in next experiments to characterize its role in plant immunity. Of the two isoforms of XBAT35, the expression pattern of *XBAT35.1* and *XBAT35.2* in *Arabidopsis* plants differed after flg22 treatment (Fig. 2b and 2c). The expression level of *XBAT35.1* in flg22-treated plants was comparable to that in plants treated with water at the time points investigated (Fig. 2b), whereas the expression of *XBAT35.2* was dramatically induced by flg22 at 60 and 90 min after treatment (Fig. 2c). We also generated stable, homozygous *Arabidopsis* transgenic lines that constitutively overexpress FLAG-tagged XBAT35.1 or XBAT35.2 and tested growth of *Pst* DC3000 on these plants. The pathogen growth on *XBAT35.1*-overexpressing plants was comparable to that on Col-0 plants (Fig. 2d). However, overexpression of *XBAT35.2* rendered the *Arabidopsis* plants more resistant to *Pst* DC3000 (Fig. 2e). These results together support the notion that XBAT35.2 plays a positive role but XBAT35.1 and XBAT34 are not involved in plant immunity against the bacterial pathogen *Pst* DC3000, which is consistent with a previous report that suggests XBAT35.2 but not XBAT35.1 is likely involved in plant immunity (Liu et al., 2017).

**Figure 2.**
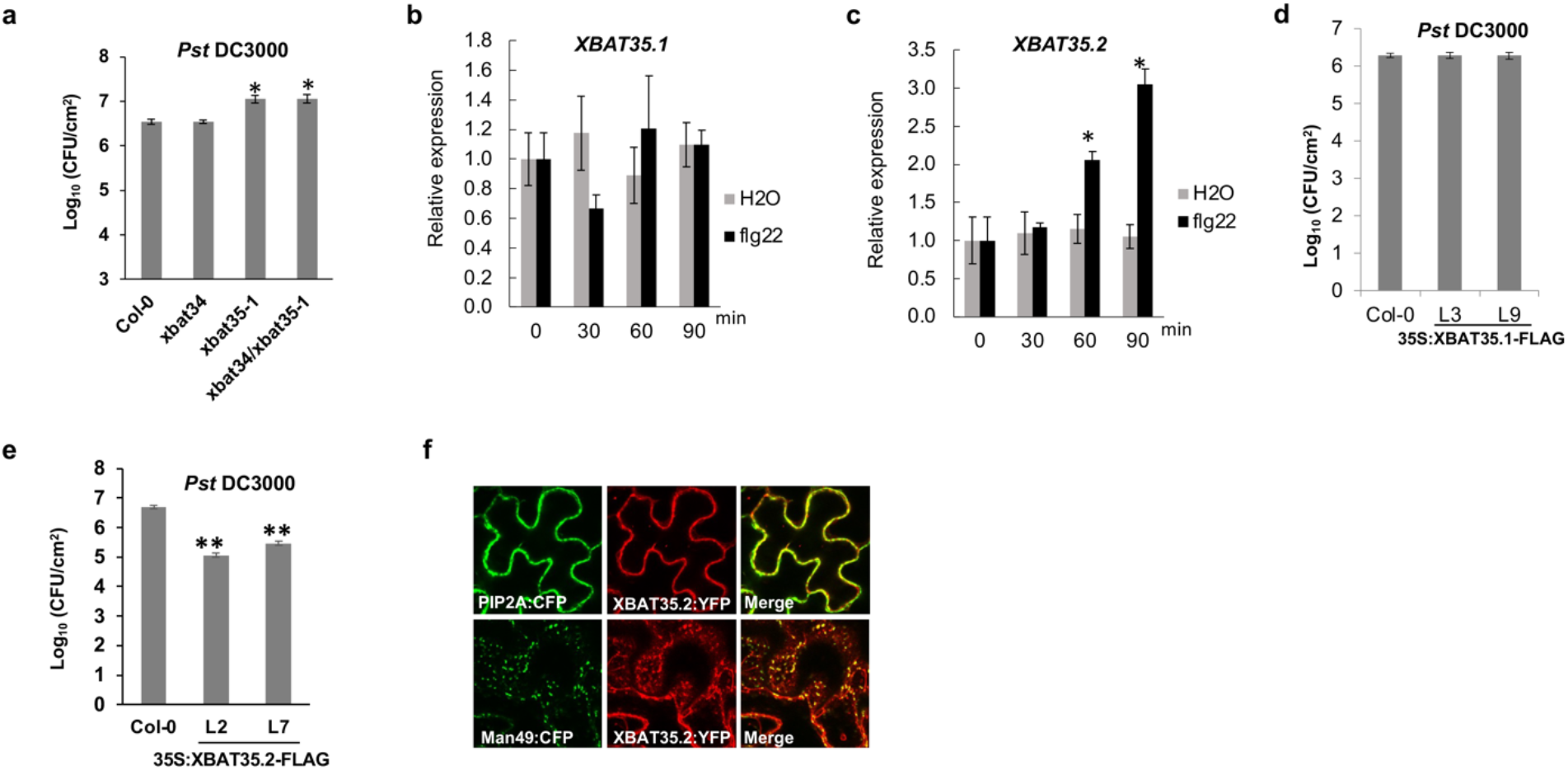
XBAT35.2 but not XBAT35.1 and XBAT34 is required for plant immunity against *Pst* DC3000 and XBAT35.2 is localized to both the Golgi body and the plasma membrane (PM). **(a), (d)** and **(e)** Growth of *Pst* DC3000 on leaves of Col-0 and the *xbat34*, *xbat35-1*, and *xbat34/35-1* mutant lines **(a)**, *XBAT35.1* overexpression **(d)**, and *XBAT35.2* overexpression **(e)** transgenic *Arabidopsis* plants. Four-week-old plants were inoculated with *Pst* DC3000 (OD_600_=0.2), and the bacterial growth was examined at 48 hours post inoculation (hpi). Error bars represent the standard deviation (SD) of 9 individual plants of each genotype. *: P< 0.05, **: P< 0.01. Experiments were repeated twice for **(d)** and three times for **(a)** and **(e)** with similar results. **(b)** and **(c)** Relative gene expression of *XBAT35.1* **(b)** and *XBAT35.2* **(c)** upon 1uM flg22 treatment, measured by qRT-PCR. Error bars represent the SD of 3 independent biological replicates. * P<0.05. Experiments were repeated three times for **(b)** and **(c)** with similar results. **(f)** Subcellular localization of XBAT35.2 in tobacco leaf epidermal cells. The YFP signal is pseudo-colored in red, and CFP signals are shown in green. PIP2A:CFP and Man49:CFP are PM and Golgi-specific marker, respectively (Nelson et al., 2007). White bar, 20 micrometers (μm). This experiment was repeated three times.

### XBAT35.2 is localized to both Golgi and PM

Live cell imaging using tobacco (*Nicotiana benthamiana*) leaf and *Arabidopsis* protoplasts showed that the XBAT35.2-YFP signal overlapped with both the Golgi (Man49-CFP) and the PM (PIP2A-CFP) marker signal (Fig. 2f and Supplementary Fig. 6a) (Nelson et al., 2007), suggesting XBAT35.2 is localized to both the Golgi body and PM. By contrast, XBAT35.1 was targeted to the nucleus (Supplementary Fig. 6b), which is consistent with a previous study (Carvalho et al., 2012). The corresponding tomato isoform XBSL35.2 also displayed localizations to both the Golgi and the PM (Supplementary Fig. 6c), further supporting the notion that XBAT35 and XBSL35 are orthologs and their roles in plant immunity might be conserved. Interestingly, XBAT34 was shown to also localize to both the PM and Golgi (Supplementary Fig. 6d). The localization of XBAT34, XBAT35, and XBSL35 to the PM is in line with the fact that they all have the consensus sequence M-G-X_3_-S-K at the N end for N-myristoyl transferase recognition, which usually suggests PM localization (Resh, 1999).

### Flg22 treatment enhances the interaction of XBAT35.2 with FLS2 and BAK1 but reduces that with BIK1

The localization of XBAT35.2 to the PM and the result that *XBAT35.2* is induced by flg22 raise the possibility that XBAT35.2 is involved in plant immunity likely through FLS2-associated processes. To test this hypothesis, we first conducted BiFC assay to investigate whether XBAT35.2 interacts with FLS2 and two closely associated protein kinases BAK1 and BIK1. For comparison, XBAT34 and XBAT35.1 were included in the assay and the GUS was used as negative control. Although XBAT34 is localized at the PM (Supplementary Fig. 6d), no YFP signal was observed for any of the combinations between XBAT34-cYFP and FLS2-nYFP, BAK1-nYFP, or BIK1-nYFP, respectively (Fig. 3a). XBAT35.1-cYFP also showed no interactions with FLS2-nYFP and BAK1-nYFP but interacted with BIK1-nYFP in the nucleus (Fig. 3a), which is consistent with that XBAT35.1 is nucleus-localized (Supplementary Fig. 6b) and BIK1 can also localize to the nucleus in addition to the PM (Lal et al., 2018). By contrast, each of the combinations of XBAT35.2-cYFP with FLS2-nYFP, BAK1-nYFP, and BIK1-nYFP, respectively produced strong YFP signal at the PM (Fig. 3a). To verify PM is the place where interaction of XBA35.2 with FLS2, BAK1, and BIK1 occurs, we added the PM-localization marker PIP2A-CFP into the BiFC assay (Nelson et al., 2007). Confocal images showed that both the CFP signal from the PM-specific marker and the YFP signals generated from interaction between XBAT35.2-cYFP and FLS2-nYFP, BAK1-nYFP and BIK1-nYFP, respectively overlapped well, indicating the interactions indeed occurred at the PM (Fig. 3b). In consistence, when an endoplasmic reticulum (ER)-specific marker (ER::CFP) (Nelson et al., 2007) was added to the BiFC assay, no overlap of CFP signal with the YFP signal from interaction of XBAT35.2-cYFP with FLS2-nYFP, BAK1-nYFP, and BIK1-nYFP was detected (Supplementary Fig. 7).

**Figure 3.**
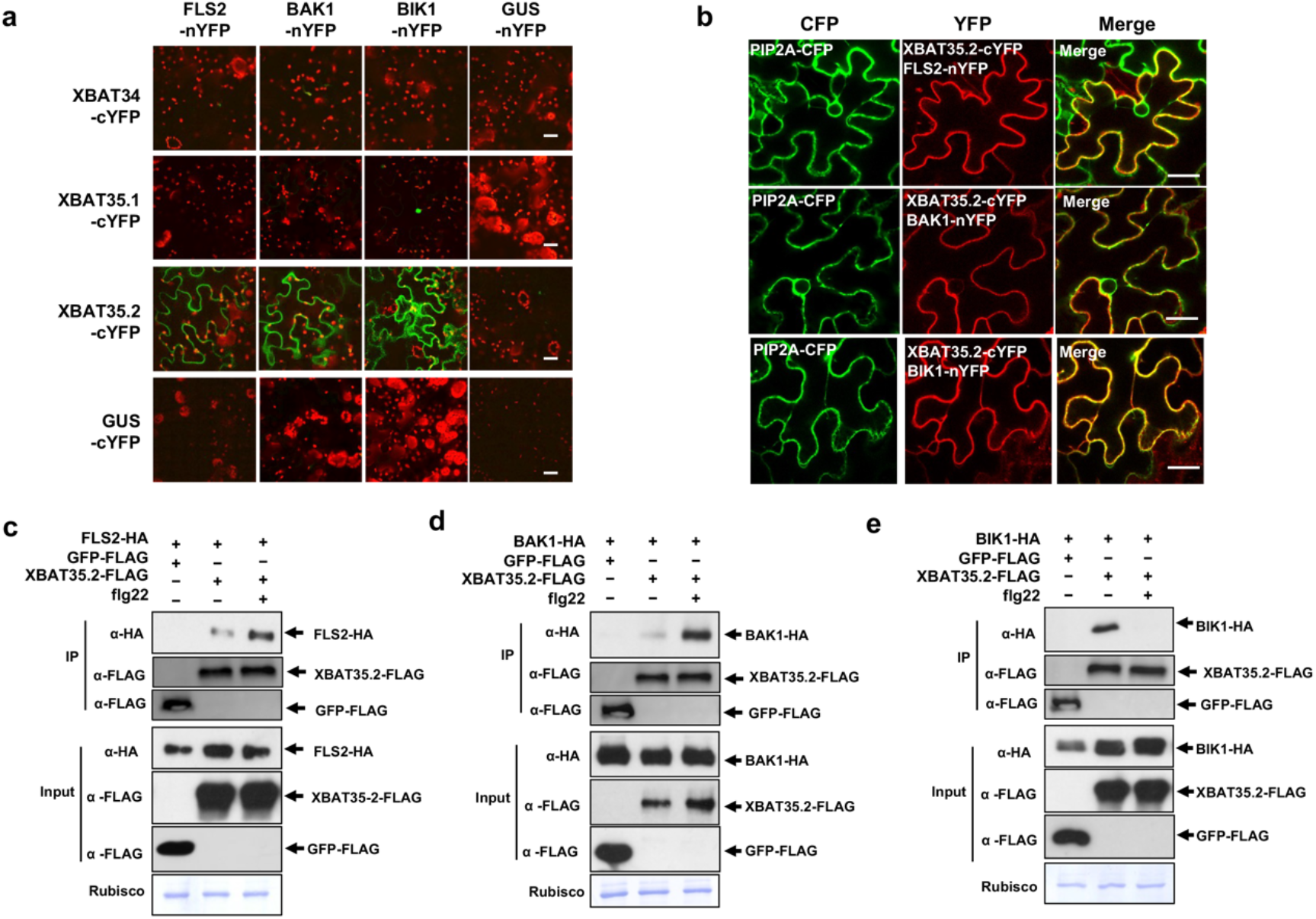
XBAT35.2 interacts with FLS2, BAK1, and BIK1 at the PM. **(a)** BiFC analysis of the interaction of XBAT34, XBAT35.1 and XBAT35.2 with FLS2, BAK1, and BIK1 in tobacco leaf epidermal cells. YFP signal (pseudo-colored in green) indicates interaction between XBAT35.2 and FLS2, BAK1 and BIK1. The YFP signal was detected 48 h after infiltration. GUS was used as negative control. White bar, 20 μm. The experiment was repeated twice with similar results. **(b)** The interaction of XBAT35.2 with FLS2, BAK1, and BIK1 occurs at the PM as indicated by PM marker PIP2A-CFP (Nelson et al., 2007) in tobacco leaf epidermal cells. The YFP signal was detected 48 h after infiltration. White bar, 20 μm. Experiment was repeated twice with similar results. **(c), (d)** and **(e)** Co-IP assays in *Arabidopsis* protoplasts confirm the interaction of XBAT35.2 with FLS2, BAK1, and BIK1. The protoplasts were co-transformed with XBAT35.2-FLAG or GFP-FLAG (as control) and FLS2-HA (**c**), BAK1-HA (**d**), and BIK1-HA (**e**). Immunoprecipitation was carried out using beads containing anti-FLAG antibody. Immunoprecipitated samples were analyzed by immunoblotting using anti-HA antibody. The amount of input proteins used for Co-IP and the bait proteins were also detected by immunoblotting using appropriate antibodies. For flg22 treatment, protoplasts were stimulated with 1 μM flg22 for 30 min before lysis. Experiments were repeated twice for **(d)**, **(e)** and three times for **(c)** with similar results.

To further confirm the interaction of XBAT35.2 with FLS2, BAK1 and BIK1, co-immunoprecipitation (Co-IP) assay was performed. XBAT35.2 could be co-immunoprecipitated with FLS2, BAK1, and BIK1 when the corresponding proteins were expressed either in *Arabidopsis* protoplasts or tobacco leaves whereas the negative control GFP did not (Fig. 3c-e, Supplementary Fig. 8a and 8b). Importantly, the association of XBAT35.2 with FLS2 and BAK1 increased upon flg22 treatment (Fig. 3c and 3d and Supplementary Fig. 8b). By contrast, flg22 elicitation significantly reduced the association of XBAT35.2 with BIK1 (Fig. 3e), which is reminiscent of the ligand-induced monoubiquitination and subsequent release of BIK1 from the FLS2-BAK1 complex (Ma et al., 2020).

To find out which region of XBAT35.2 is responsible for interaction with FLS2, BAK1 and BIK1, we created two truncated versions of XBAT35.2, XBAT35.2^ΔANK^, and XBAT35.2^ΔRING^ for BiFC assay (Supplementary Fig. 9a). As shown in Supplementary Fig. 9b, deletion of the ANK repeats domain compromised the interaction of XBAT35.2 with FLS2, BAK1 and BIK1. However, the truncated version XBAT35-2^ΔRING^ in which the ANK repeats domain remains intact interacted with FLS2, BAK1, and BIK1 at similar level to that of the full-length XBAT35.2 (Supplementary Fig. 9b). These findings indicated that the ANK repeat domain rather than the RING domain is essential for the interactions of XBAT35.2 with FLS2, BAK1, and BIK1.

### XBAT35.2 stabilizes FLS2 and increases host immunity to bacterial pathogens

The results that XBAT35.2 is an active E3 ligase and interacts with FLS2 prompted us to evaluate whether XBAT35.2 functions in plant immunity through modulating the abundance of FLS2 by ubiquitination. To this end, we first tested the protein level of FLS2 in Col-0, *xbat35-1* mutant, and *XBAT35.2*-overexpressing (*XBAT35.2*-OEX) plants using anti-FLS2 antibodies. The specificity of the anti-FLS2 antibody was confirmed by immunoblotting that includes the fls2 null mutant (Supplementary Fig. 10). As shown in Fig. 4a, 4b and supplementary Fig. 10, without flg22 treatment, the FLS2 protein level in *xbat35-1* is comparable to that in Col-0 whereas the abundance of FLS2 is increased in two independent *XBAT35.2*-OEX *Arabidopsis* lines. FLS2 has been shown to undergo flg22-induced endocytosis and subsequent degradation (Robatzek et al., 2006; Beck et al., 2012). Consistently, the FLS2 protein level in Col-0 and the *xbat35-1* mutant line was notably reduced 60 minutes after flg22 treatment (Fig. 4a and 4c). Nevertheless, the decrease of FLS2 in *xbat35-1* is more significant compared to that in Col-0 (Fig. 4a). By contrast, the abundance of the FLS2 protein remained at a much higher level in *XBAT35.2*-OEX lines than that in Col-0 after flg22 treatment, with less flg22-induced degradation compared to that in Col-0 (Fig. 4c), which is in consistence with the increased and reduced *Pst* DC3000 growth in the *xbat35-1* mutant and *XBAT35.2*-OEX lines, respectively (Fig. 2a, 2e). By contrast, the abundance of the BAK1 protein was not affected by overexpression of *XBAT35.2* (Fig. 4b), indicating the effect of XBAT35.2 on FLS2 abundance is specific. To ensure equal loading of samples in these experiments, we examined the Actin protein level in the samples in addition to checking the abundance of the Rubisco protein. Both immunoblotting analysis of Actin and staining of the Rubisco confirmed equal loading. The consistence of the results by the two methods validated the use of the abundance of Rubisco for checking equal sample loading, which was further supported in subsequent experiments (Fig. 7b). To investigate whether the increased accumulation of FLS2 is caused by change of the *FLS2* gene transcription in the *XBAT35.2*-OEX plants, we quantified the relative transcript level of *FLS2* in Col-0 and *XBAT35.2*-OEX plants with and without flg22 treatment. The expression level of *FLS2* in *XBAT35.2*-OEX plants was comparable to that in the Col-0 regardless of flg22 treatment (Fig. 4d), suggesting XBAT35.2 promotes the FLS2 accumulation at the protein but not mRNA level. In line with the increased accumulation of FLS2 in *XBAT35.2*-OEX lines, seedling growth inhibition assay indicated that the *XBAT35.2*-OEX lines are more sensitive to flg22-induced growth inhibition (Fig. 4e-f) (Gómez-Gómez and Boller, 2000). We then tested whether XBAT35.2 can ubiquitinate FLS2. The *in vitro* ubiquitination assay indicated that the FLS2 cytosolic domain (CD) was not ubiquitinated by XBAT35.2, whereas XBAT35.2 showed strong auto-ubiquitination and the positive control *Arabidopsis* PUB12 showed strong ubiquitination of FLS2CD as previously reported (Lu et al., 2011) (Fig. 4g). Taken together, these findings suggest that XBAT35.2 affects the FLS2 protein level in *Arabidopsis* plants not by targeting FLS2 for ubiquitination, but instead, likely through third-party components.

**Figure 4.**
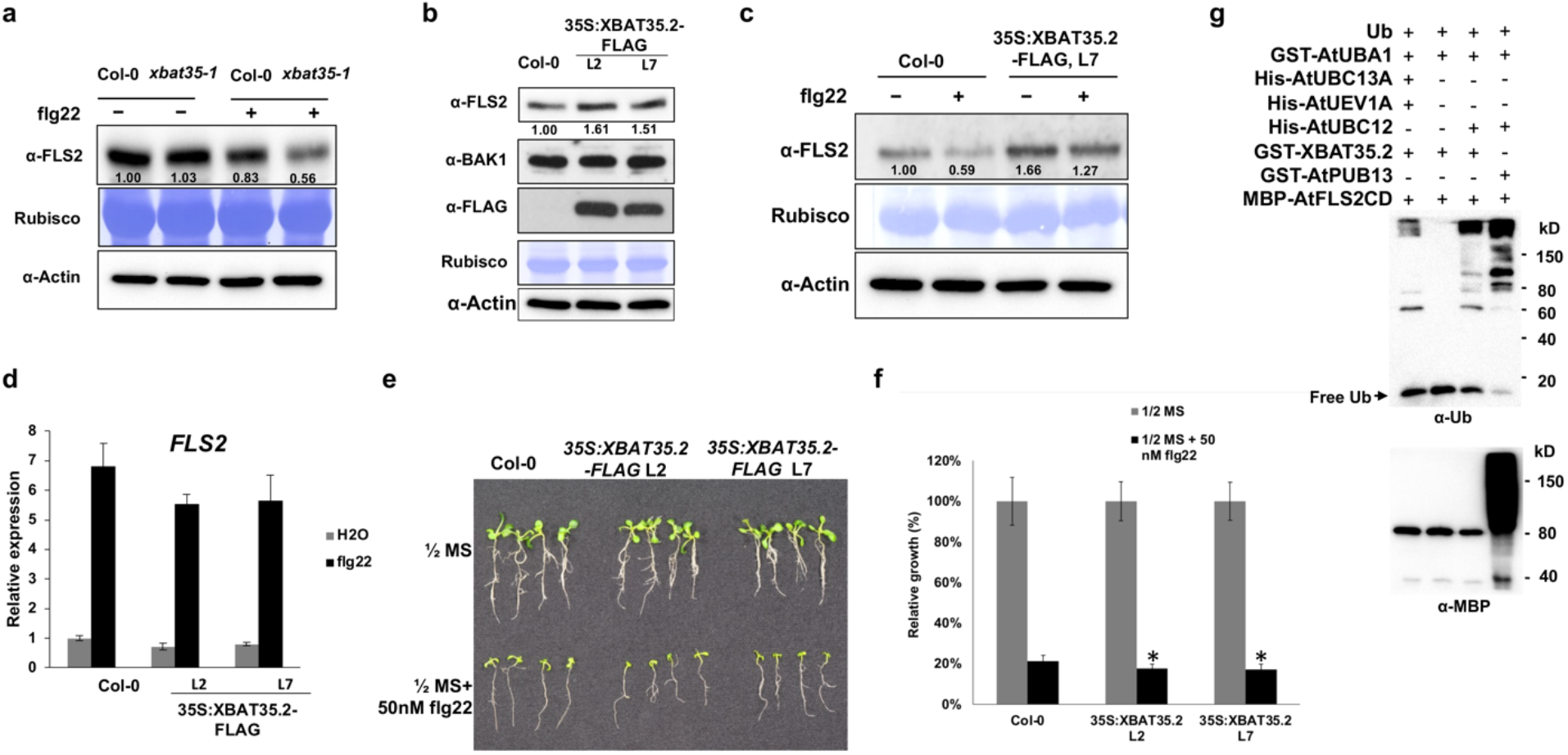
XBAT35.2 stabilizes FLS2 and enhances plant immunity to *Pst* DC3000. **(a)** The protein level of FLS2 in Col-0 and the *xbat35-1* mutant with or without flg22 treatment. Ten-day-old seedlings grown on ^1^/_2_ MS media were transferred into liquid ^1^/_2_ MS with or without 1μM flg22 for 60 min, and the total proteins were then extracted and followed by immunoblotting using anti-FLS2 antibody. The numbers below the bands denote the relative abundance of the FLS2 protein with the normalized value in untreated Col-0 plants being set as 1. Both staining of the Rubisco and anti-Actin immunoblot indicated equal loading of the samples. The experiment was repeated twice with similar results. **(b)** and **(c)** The protein level of FLS2 in Col-0 and *XBAT35.2*-overexpressing lines with or without flg22 treatment. The total proteins were extracted from 10-day-old seedlings grown on ^1^/_2_ MS media and followed by immunoblotting using corresponding antibody. The numbers below the bands denote the relative abundance of the FLS2 protein, with the normalized value in untreated Col-0 plants being set as 1. Both staining of the Rubisco and anti-Actin immunoblot indicated equal loading of the samples. The experiments were repeated three times with similar results. **(d)** The relative expression of *FLS2* in Col-0 and *XBAT35.2*-overexpressing lines with flg22 or H_2_O (as control) treatment. The experiments were repeated three times with similar results. **(e)** *XBAT35.2*-overexpressing lines are more sensitive than Col-0 to 50 nM flg22-induced inhibition of seedling growth. Photo was taken 10 days after the four-day-old seedlings were treated with flg22. The experiment was repeated twice with similar results. **(f)** Statistical analysis of the relative growth of Col-0 and *XBAT35.2*-overexpressing lines in **(e)**. Error bars represent the SD of 6 independent samples. *: P< 0.05. The experiment was repeated twice with similar results. **(g)** XBAT35.2 does not ubiquitinate FLS2 in *in vitro* ubiquitination assay. The *Arabidopsis* E3 AtPUB13 was used as positive control. Anti-Ub antibody was used to detect the auto-ubiquitination of XBAT35.2 and AtPUB13 whereas anti-MBP antibody was used to detect the ubiquitination of MBP-tagged FLS2 cytosolic domain (FLS2CD). The experiment was repeated twice with similar results.

### XBAT35.2 interacts with VPS37-1 and VPS28-2 *in planta*

The protein level of FLS2 has been shown to be modulated by both PUB12/13-mediated ubiquitination and endocytosis-mediated recycling and degradation (Robatzek et al., 2006; Lu et al., 2011; Beck et al., 2012). Because XBAT35.2 does not ubiquitinate FLS2 yet positively regulates its abundance (Fig. 4a-c), we hypothesize that XBAT35.2 stabilizes FLS2 by disturbing the endocytosis-mediated breakdown of FLS2. We thus pay special attention to components involved in the endocytosis and ensuing trafficking/sorting of FLS2 when screening for putative targets of XBAT35.2. To date, the dynamin-related proteins DRP1A and DRP2B, and ECSRT-I components VPS28-2 and VPS37-1 have been demonstrated to be implicated in endocytosis and endosomal trafficking/sorting of FLS2 (Spallek et al., 2013; Smith et al., 2014; Ekanayake et al., 2021). Accordingly, we tested these proteins.

We first investigated the interaction of XBAT35.2 with DRP1A, DRP2B, VPS28-2 and VPS37-1 using BiFC assay. As shown in Fig. 5a, confocal microscopy showed reconstituted YFP fluorophore between VPS37-1-cYFP and XBAT35.2-nYFP, implying that the two proteins interact with each other. Strong interactions were detected between VPS37-1-cYFP and the E3-ligase null mutant XBAT35.2^AA^-nYFP (Liu et al., 2017), suggesting that XBAT35.2 could interact with VPS37-1 independent of its E3 activity. To test this, we employed the truncations of XBAT35.2, XBAT35.2^ΔANK^, and XBAT35.2^ΔRING^ for the BiFC assay. As shown in Supplementary Fig. 11a, VPS37-1-nYFP showed interaction with XBAT35.2^ΔRING^-cYFP, but not with XBAT35.2^ΔANK^- cYFP, suggesting the ANK repeats domain of XBAT35.2 is essential for interaction with VPS37-1 and corroborating the notion that ANK repeats domain facilitates protein-protein interactions (Kane and Spratt, 2021). Like VPS37-1, VPS28-2 was also observed to interact with XBAT35.2 in an ANK domain-dependent manner (Supplementary Fig. 11b). No fluorescence signals were detected when GUS-cYFP or GUS-nYFP (as negative control) was co-expressed with VPS37-1 or XBAT35.2 with corresponding split YFP (Fig. 5a and 5b). Like previously reported (Spallek et al., 2013), we also observed the interaction of FLS2 with VPS37-1 and VPS28-2 (Fig. 5a and 5b). Importantly, we didn’t observe YFP signal when DRP1A and DRP2B was co-expressed with XBAT35.2 in the assay (Fig. 5c). By contrast, strong YFP signals were detected when DRP1A-cYFP or DRP2B-cYFP was co-expressed with FLS2-nYFP, which is consistent with previous reports (Smith et al., 2014; Ekanayake et al., 2021).

**Figure 5.**
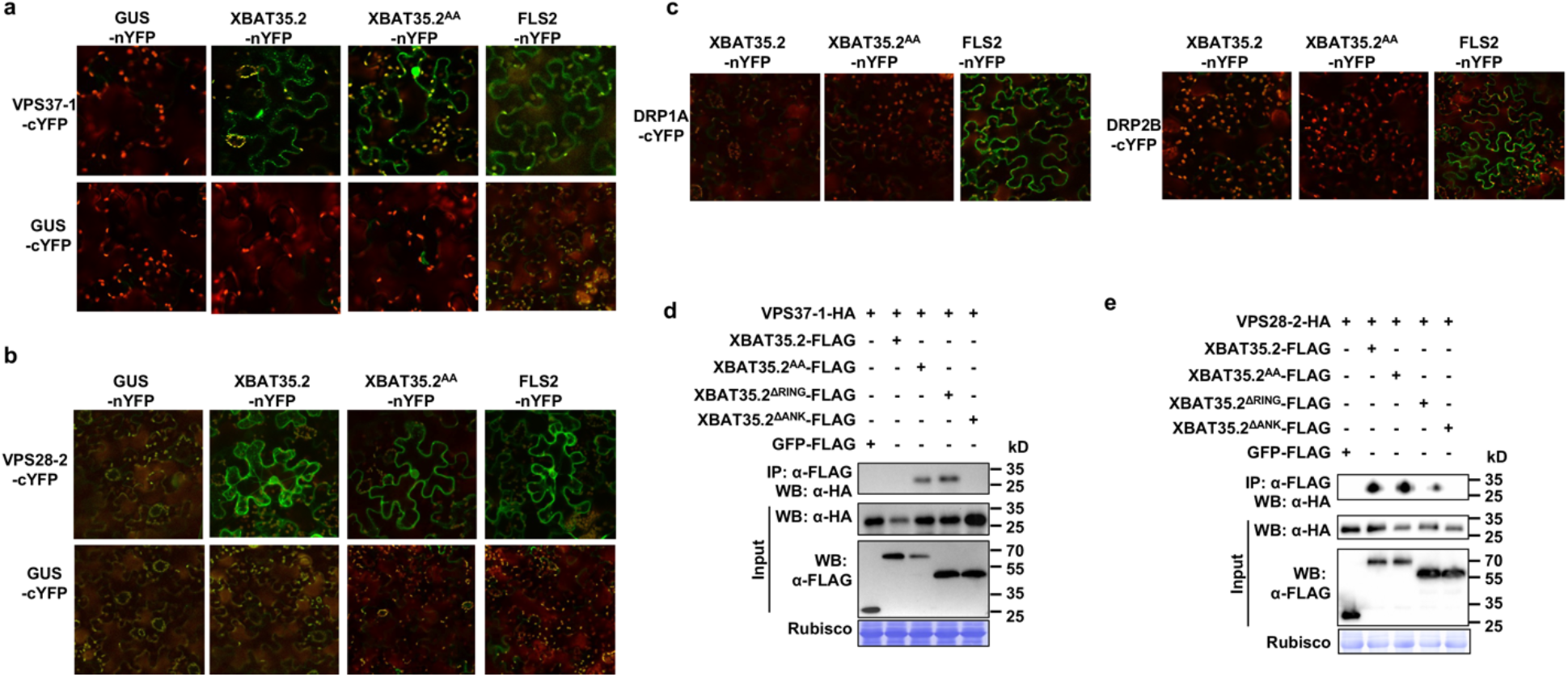
XBAT35.2 specifically interacts with VPS28-2 and VPS37-1 *in vivo*. **(a)** and **(b)** BiFC analysis of the interaction of cYFP-fused VPS37-1 (**a**) and VPS28-2 (**b**) with nYFP-fused XBAT35.2, its truncation mutants (XBAT35.2^ΔANK^, XBAT35.2^ΔRING^) and E3 ligase activity null mutant (XBAT35.2^AA^) in tobacco leaf epidermal cells. YFP signal (pseudo-colored in green) indicates interaction between the two proteins tested. The tobacco leaves were collected and examined on the third day after Agro-infiltration. GUS and FLS2 were used as negative and positive control, respectively. The experiments were repeated twice with similar results. **(c)** XBAT35.2 does not interact with DPP1A and DRP2B in BiFC assay. YFP signal (pseudo-colored in green) indicates interaction between the two proteins tested. FLS2 was used as positive control. The experiment was repeated twice with similar results. **(d)** and **(e)** Co-IP assay confirms the interaction of XBAT35.2 with VPS37-1 (**d**) and VPS28-2 (**e**) *in vivo*. *A. tumefaciens* cells harboring the constructs that express VPS37-1-HA or VPS28-2-HA and XBAT35.2-FLAG, XBAT35.2^ΔANK^-FLAG, XBAT35.2^ΔRING^-FLAG, or XBAT35.2^AA^-FLAG were co-infiltrated into tobacco leaves. Total protein was extracted from leaves on the third day after Agro-infiltration. Co-immunoprecipitation was carried out with anti-FLAG antibody, and the immuno-precipitated samples were analyzed by immunoblotting using anti-HA antibody. GFP-FLAG was used as negative control. The experiments were repeated twice with similar results.

The interactions of XBAT35.2 with VPS37-1 and VPS28-2 were further confirmed by Co-IP assay. As shown in Fig. 5d, VPS37-1-HA could be co-immunoprecipitated by XBAT35.2^AA^-FLAG and XBAT35.2^ΔRING^-FLAG, but not by XBAT35.2^ΔANK^-FLAG, corroborating the conclusion drawn from the BiFC assay (Fig. 5a) that the ANK domain of XBAT35.2 is required for interaction with VPS37-1. However, VPS37-1-HA was not co-immunoprecipitated by wild type XBAT35.2-FLAG (Fig. 5d), apparently because that association of enriched XBAT35.2 as an active E3 ligase with VPS37-1 during Co-IP led to the ubiquitination and degradation of VPS37-1 by proteasome. Indeed, application of MG132 restored the co-precipitation of VPS37-1 with XBAT35.2 (Supplementary Fig. 11c). The co-immunoprecipitation of VPS28-2-HA with XBAT35.2-FLAG, XBAT35.2^AA^-FLAG, and XBAT35.2^ΔRING^-FLAG was also detected by the Co-IP assay (Fig. 5e), which is in consistence with the result shown in Fig. 5b. The Co-IP of XBAT35.2 with VPS28-2 suggests that it likely does not target VPS28-2 for degradation *in planta*.

### XBAT35.2 ubiquitinates VPS28-2 and VPS37-1 through distinct linkage types of ubiquitin chains

We next examined whether XBAT35.2 can ubiquitinate VPS37-1 and VPS28-2 using *in vitro* ubiquitination assay. As shown in the lower panel of Supplementary Fig. 12a, in the presence of E1 (GST-AtUBA1), E2 (GST-AtUBC12), ubiquitin, and other essential factors, MBP-VPS37-1 could be conjugated with ubiquitin chains in the presence of GST-XBAT35.2, but not when XBAT35.2 was absent. By contrast, the MBP tag along as negative control was not ubiquitinated by XBAT35.2 even though strong autoubiquitination was detected for XBAT35.2 (Supplementary Fig. 12a, upper panel). Like a previous report that only faint ubiquitination of VPS23A by XBAT35 was detected in *in vitro* ubiquitination (Yu et al., 2020), we detected weak signal in the ubiquitination VPS37 by XBAT35.2 *in vitro*. To find out the type of ubiquitin linkage conjugated to VPS37-1 protein, we utilized ubiquitin variants-UbK48R where the K48 amino acid residue is mutated to R and UbK48only in which all seven Lys amino acid residues except for K48 are mutated to R and revealed VPS37-1 is modified with K48-linked ubiquitin chains (Fig. 6a). Ubiquitin E2 AtUBC13A and its cofactor AtUEV1a that direct exclusively K63-linked ubiquitination as well as ubiquitin variants UbK63R and UbK63only were used in a parallel *in vitro* ubiquitination assay. As shown in Supplementary Fig. 12b (lanes 1, 3, 5, and 7), while we detected XBAT35.2 autoubiquitination, no ubiquitination of VPS37-1 was detected, indicating VPS37-1 cannot be modified by XBAT35.2 through K63-linked ubiquitination. Importantly, VPS28-2 was found to be ubiquitinated by XBAT35.2 by K63-linked ubiquitination when AtUBC13A and its cofactor AtUEV1a were employed (Fig. 6b). By contrast, when the E2 UBC12 that directs K48-linked ubiquitination was used, VPS28-2 was not ubiquitinated by XBAT35.2. To further explore the effects of XBAT35.2 on VPS37-1 and VPS28-2, we also created stable transgenic lines by transformation of *35S:VPS37-1-HA* or *35S:VPS28-2-HA* constructs into Col-0, the *xbat35-1* mutant and *35S:XBAT35.2-FLAG Arabidopsis* plants. Immunoblotting indicated that the ubiquitination of VPB37-1 and VPS28-2 is much stronger in the *35S:XBAT35.2-FLAG Arabidopsis* plants whereas more significantly reduced in the *xbat35-1* mutant than that in Col-0 (Supplementary Fig. 13), indicating that VPS37-1 and VPS28-2 are ubiquitinated by XBAT35.2 *in planta*.

**Figure 6.**
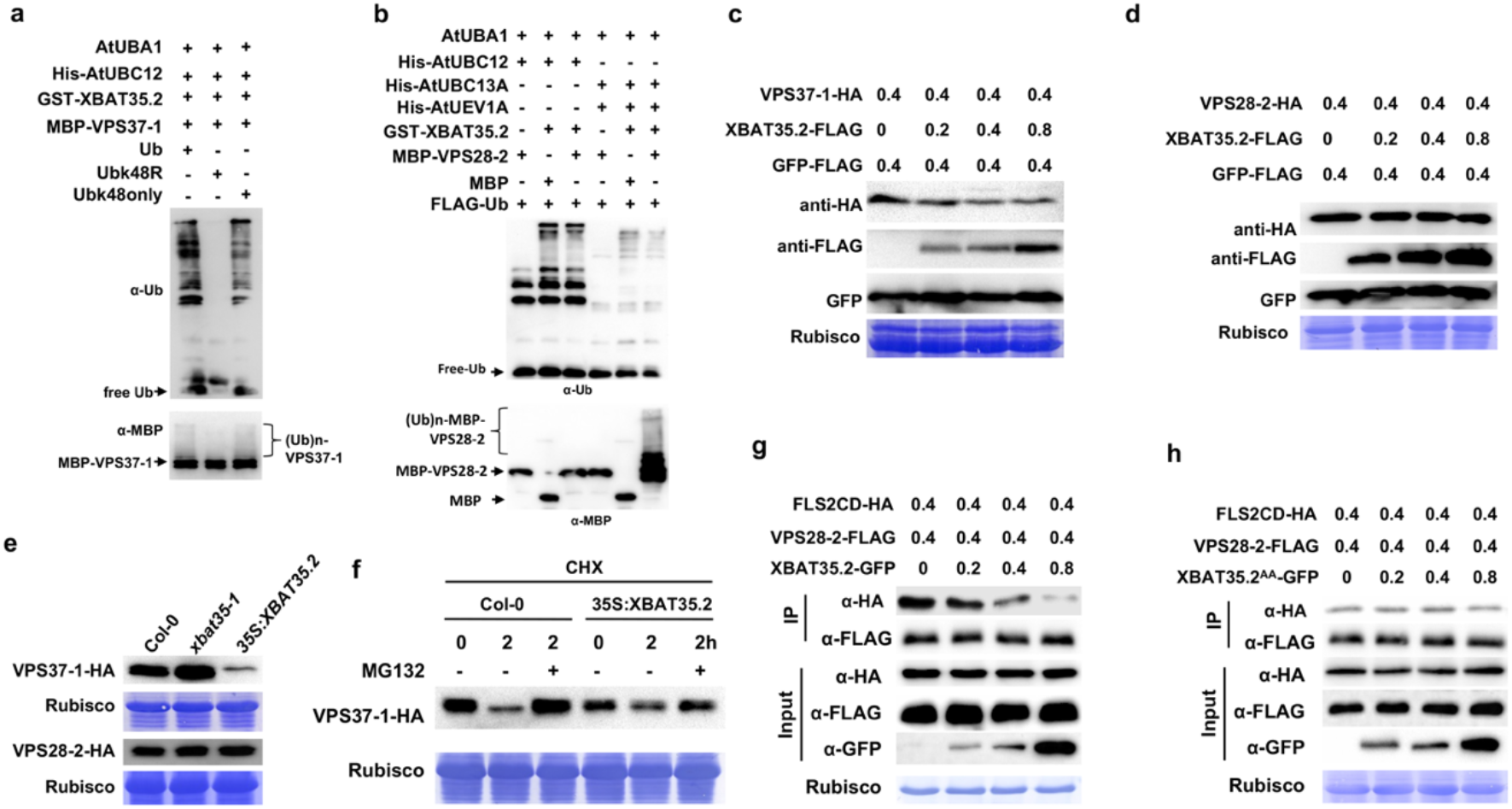
XBAT35.2 directs ubiquitination of VPS37-1 and VPS28-2 using distinct ubiquitin chains. **(a)** XBAT35.2 works with the ubiquitin E2 UBC12 to ubiquitinate VPS37-1 with K48-linked ubiquitin chains. Upper panel: XBAT35.2 autoubiquitination detected by anti-ubiquitin antibody. Lower panel: The high molecular weight smear marks ubiquitination of VPS37-1 [(Ub)n-VPS37-1]. MBP was used as negative control. This experiment was repeated twice with similar results. **(b)** XBAT35.2 works with AtUBC13A/AtUEV1a to ubiquitinate VPS28-2 *in vitro* through K63-linked ubiquitin chains. Upper panel: XBAT35.2 autoubiquitination detected by anti-ubiquitin antibody denotes E3 ligase activity of XBAT35.2. Lower panel: The high molecular weight smear represents ubiquitination of VPS28-2 [(Ub)n-VPS28-2]. MBP was used as negative control. This experiment was repeated twice with similar results. **(c)**, **(e)** and **(f)** XBAT35.2 promotes the degradation of the VPS37-1 protein in 26S proteasome-dependent manner. *A. tumefaciens* carrying *35S:VPS37-1-HA* constructs were co-infiltrated into tobacco leaves with increasing amount of *A. tumefaciens* cells that harbor the *35S:XBAT35.2-FLAG* constructs, and GFP was used as internal expression control **(c)**. The null mutation of XBAT35 blocks VPS37-1 protein degradation whereas overexpression of XBAT35.2 significantly enhances the degradation of VPS37-1 (**e**). However, neither disruption nor overexpression of XBAT35.2 affects the protein level of VPS28-2. The transgenic seedlings where VPS28-2 or VPS37-1-HA is expressed in the genetic background of Col-0, *xbat35-1*, and *35S:XBAT35.2-FLAG*, respectively, were grown in soil for 4 weeks, followed by total proteins extraction and immunoblotting analysis **(e)**. MG132 treatment significantly inhibits degradation of VPS37-1-HA protein **(f)**. Ten-day-old seedlings of *Arabidopsis* transgenic lines where VPS37-1-HA is expressed in the genetic background of Col-0 and *35S:XBAT35.2-FLAG* were treated with 25 μM CHX or 25 μM CHX and 75 μM MG132 as indicated for 2 h before sampling. The experiment for **(c)** was repeated three times and for **(e)** was repeated twice with similar results. **(d)** XBAT35.2 does not cause degradation of VPS28-2. *A. tumefaciens* carrying the *35S:VPS28-2-HA* construct were co-infiltrated into tobacco leaves with increasing amount of *A. tumefaciens* cells that harbor the *35S:XBAT35.2*-*FLAG* construct. GFP-FLAG was used as internal expression control. The experiment was repeated three times with similar results. **(g)** and **(h)** XBAT35.2 promotes the disassociation of VPS28-2 with FLS2, which is dependent on its E3 ligase activity. The same amount of *A. tumefaciens* cells transformed with *35S:VPS28-2-FLAG* and *35S:FLS2CD-HA* were co-infiltrated into tobacco leaves with increasing amount of *A. tumefaciens* cells that express XBAT35.2-GFP (**f**) or the E3 ligase activity null mutant XBAT35.2^AA^-GFP (**g**). Samples were collected 48 h after Agro-infiltration and co-immunoprecipitation was carried out with anti-FLAG antibody. The immunoprecipitated samples were analyzed by immunoblotting using anti-HA antibody. The experiments were repeated twice with similar results.

### XBAT35.2-mediated ubiquitination results in degradation of VPS37-1 and significantly reduced interaction of VPS28-2 with FLS2

K48-linked polyubiquitin chains usually serve as a signal for protein degradation by the 26S proteasome whereas K63-linked ubiquitination generally marks target proteins for non-proteolytic, regulatory outcomes. To explore whether XBAT35.2 promotes degradation of VPS37-1 and VPS28-2, we performed degradation assays in tobacco leaves and our results indicated that VPS37-1 was gradually reduced as the amount of XBAT35.2 increased (Fig. 6c). By contrast, the protein level of VPS28-2 was not affected by XBAT35.2, regardless of the amount of co-expressed XBAT35.2 (Fig. 6d). Consistent with the previous report (Yu et al., 2020), a parallel degradation assay experiment found that VPS23A was gradually reduced as the amount of XBAT35.2 increased (Supplementary Fig. 14), validating our degradation assays performed on tobacco leaves. Immunoblotting using the stable transgenic lines where *35S:VPS37-1-HA* or *35S:VPS28-2-HA* is expressed in Col-0, the *xbat35-1* mutant and *35S:XBAT35.2-FLAG Arabidopsis* plants showed that there was a higher accumulation of VPS37-1 in *xbst35-1* mutant compared with that in Col-0, whereas the protein level of VPS37-1-HA decreased sharply in *35S:XBAT35.2-FLAG* lines (Fig. 6e), suggesting that XBAT35.2 targets VPS37-1 for proteolysis *in vivo*. By contrast, the Col-0, *xbat35-1,* and *35S:XBAT35.2-FLAG Arabidopsis* plants displayed comparable level of VPS28-2-HA (Fig. 6e). To investigate whether the XBAT35.2-mediated VPS37-1 degradation was 26S proteasome-dependent, we did a cycloheximide (CHX) chase degradation assay using 2-week-old transgenic *Arabidopsis* seedlings expressing VPS37-1-HA in the background of wild type and *35S:XBAT35.2-FLAG* respectively. Given that the VPS37-1 protein level in the *35S:XBAT35.2* background is very low even under normal growth conditions, we treated these transgenic seedlings with lower concentrations of CHX. As shown in Fig. 6f, two-hour CHX treatment significantly reduced the amount of VPS37-1 in both Col-0 and *35S:XBAT35.2* plants apparently due to XBAT35.2-mediated degradation. However, the degradation of VPS37-1 was efficiently alleviated when the 26S proteasomes inhibitor MG132 was added. Worthy of notice, the overall protein level of VPS37-1 in the 35S:XBAT35.2 is lower than that in Col-0 conceivably due to XBT35.2-mediated ubiquitination and degradation. These results suggested that XBAT35 mediated degradation of VPS37-1 in a 26S proteasome-dependent manner.

K63-linked ubiquitination usually serves as non-proteolytic signal for modulating substrate proteins in activity, subcellular localization, or interaction with partners. VPS28-2 is shown to be modified by XBAT35.2 through K63-linked ubiquitination in this study (Fig. 5b) and it was shown previously to associate with FLS2 for endosomal sorting to MVB (Spallek et al., 2013). Thus, we attempted to determine whether XBAT35.2 influences the interaction of VPS28-2 with FLS2. To this end, we co-infiltrated the same amount of *Agrobacterium tumefaciens* (*A. tumefaciens*) harboring constructs that express VPS28-2-FLAG and FLS2CD-HA protein, respectively with increasing amounts of *A. tumefaciens* carrying the XBAT35.2-GFP constructs. Total proteins were extracted from leaf samples after 48 h and subject to Co-IP assays. As shown in Fig. 6g, FLS2CD-HA was co-immunoprecipitated by VPS28-2-FLAG. However, the co-immunoprecipitated FLS2CD-HA was gradually reduced as the amount of XBAT35.2 increased. By contrast, the amount of FLS2CD-HA co-immunoprecipitated with VPS28-2-FLAG was unaltered when the E3 ligase-inactive variant XBAT35.2^AA^ was used (Fig. 6h). These results indicate that XBAT35 reduced the interaction of VPS28-2 with FLS2 in an E3 ubiquitin ligase activity-dependent manner.

### VPS37-1 and VPS28-2 play redundant, negative roles in FLS2-mediated immunity by vacuolar breakdown of FLS2

The results we accumulated so far (Fig. 2a, 2c, 2e, Fig. 4a-4c, and Fig. 6) led us to hypothesize that VPS37-1 and/or VPS28-2 play a negative role in FLS2-mediated immunity, probably by promoting its turnover at the vacuole. To test this hypothesis, we identified homozygous T-DNA insertion mutant lines, *vps37-1.1* (CS804682) and *vps28-2* (SALK_040274C) (Supplementary Fig. 15a and 15b) and created double mutant line *vps28-2/vps37-1.1* by crossing. The single and double mutant plants display normal growth phenotype like the wild type plants (Supplementary Fig. 15c). We next investigated the protein level of FLS2 in these plants and found that, without flg22 treatment, the FLS2 level in single mutant *vps28-2* and *vps37-1.1* was comparable to that in Col-0. By contrast, the FLS2 protein level increased slightly in double mutant *vps28-2*/*vps37-1.1* (Fig. 7a left panel). Flg22 treatment led to significant decrease in FLS2 abundance in Col-0. However, the decrease of FLS2 in *vps37-1.1*, *vps28-2* and *vps28-2/vps37-1.1* was less than that in Col-0, with the *vps28-2/vps37-1.1* double mutant retaining a higher level of FLS2 than that in Col-0, *vps37-1.1*, and *vps28-2* (Fig. 7a right panel). We also inspected the FLS2 protein in *Arabidopsis* plants that express *35S:VPS28-2-HA* and *35S:VPS37-1-HA*, respectively. The FLS2 protein level was slightly lower in *VPS28-2-HA* or *VPS37-1-HA*-overexpressing lines than that in Col-0 without flg22 treatment (Fig. 7b left panel). Flg22 treatment induced a reduction of FLS2 abundance both in Col-0 and *VPS28-2-HA* or *VPS37-1-HA*-overexpressing line, with the overexpression of *VPS28-2* or *VPS37-1* causing much greater decline of FLS2 protein level (Fig. 7b, middle panel). These findings suggested VPS28-2 and VPS37-1 redundantly promote degradation of both nonactivated and flg22-activated FLS2 and the promotion is more significant after flg22-treatment.

**Figure 7.**
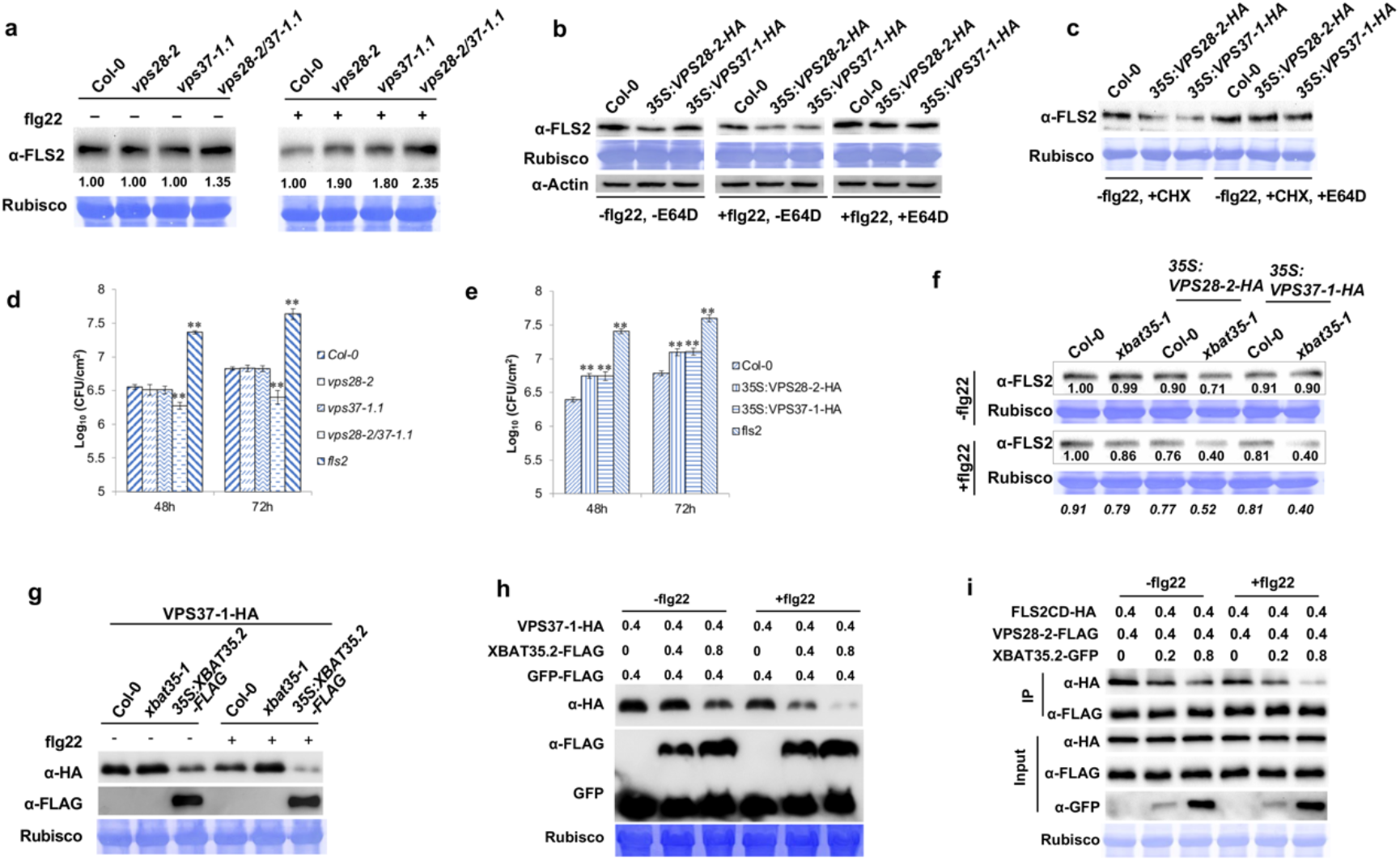
XBAT35.2 regulates FLS2-mediated plant immunity via VPS28-2 and VPS37-1 that redundantly promote vacuolar degradation of FLS2. **(a)** Loss of function in VPS28-2 and VPS37-1 redundantly inhibit flg22-induced FLS2 degradation. Ten-day-old seedlings of Col-0 and stable transgenic lines grown on ^1^/_2_ MS media were transferred into liquid ^1^/_2_ MS with or without flg22 for 1 h. The numbers below the bands denote the relative abundance of the FLS2 protein, with the normalized value in untreated Col-0 plants being set as 1. The experiment was repeated three times with similar results. **(b)** and **(c)** VPS28-2 and VPS37-1 promote vacuolar degradation of FLS2. Ten-day-old seedlings of Col-0, 35S:*VPS28-2-HA,* and 35S:*VPS37-1-HA* transgenic *Arabidopsis* plants were treated with or without flg22 and E64D (10 μg/mL) for 1 h before sampling for immunoblotting analysis **(b)**. Both staining of the Rubisco and anti-Actin immunoblot indicated equal loading of the samples. VPS28-2 and VPS37-2 promote vacuolar degradation of non-*de novo* synthesized, nonactivated FLS2 protein **(c)**. Ten-day-old seedlings of Col-0, 35S:*VPS28-2-HA*, and 35S:*VPS37-1-HA* transgenic *Arabidopsis* line were treated with 50 μM CHX or both 50 μM CHX and E64D (10 μg/mL) before sampling for immunoblotting analysis. The CHX was used to minimize the effects of *de novo* synthesized FLS2 protein on the assay. The experiments were repeated twice with similar results. **(d)** and **(e)** VPS28-2 and VPS37-1 redundantly modulate plant immunity as shown by pathogen growth assay on *Arabidopsis* plants in which *VPS28-2* and *VPS37-1* are loss-of-function (**d**) or overexpressed (**e**). The *fls2* mutant was included as control. For both (**d**) and (**e**), 4-week-old plants were inoculated with *Pst* DC3000 (OD_600_=0.2), and the bacterial growth was determined at 48 and 72 hpi. Error bars represent the SD of 9 independent samples for each genotype. **: P< 0.01. The experiments were repeated three times with similar results. **(f)** The FLS2 protein level in Col-0, *xbat35-1*, *35S:VPS28-2-HA*/Col-0, 35S*:VPS28-2-HA*/*xbat35-1*, 35S*:VPS37-1-HA*/*Col-*0, and *35S:VPS37-1-HA*/*xbat35-1* transgenic lines in the absence and presence of flg22 treatment. Ten-day-old seedlings were treated with 1 μM flg22 for 60 minutes before sampling for protein extraction. The numbers below the bands denote the relative abundance of the FLS2 protein, with the normalized value in untreated and treated Col-0 plants being set as 1. The italicized numbers at the bottom denote the ratio of ImageJ-quantified and normalized relative abundance of the FLS2 in flg22-treated sample compared to the corresponding non-flg22-treated sample for each genotype. This experiment was repeated twice with similar results. **(g)** Flg22 treatment enhances XBAT35.2-mediated degradation of VPS37-1 in *Arabidopsis*. Ten-day-old seedlings of the *Arabidopsis* transgenic line where VPS37-1-HA is expressed in the genetic background of Col-0, *xbat35-1,* and *35S:XBAT35.2-FLAG* were treated with or without flg22 for 1h before sampling for immunoblotting analysis. The experiment was repeated twice with similar results. **(h)** Flg22 treatment enhances XBAT35.2-mediated degradation of VPS37-1 in tobacco leaves. The same amount of *A. tumefaciens* cells expressing VPS37-1-HA or GFP-FLAG were co-infiltrated into tobacco leaves with increasing amount of *A. tumefaciens* cells that carry the *35S:XBAT35.2-FLAG* constructs. Tobacco leaves were sampled 3 days after Agro-infiltration. For flg22 treatment, 1uM flg22 was infiltrated into Agro-infiltrated tobacco leaves 30 min before sampling. The experiment was repeated twice with similar results. **(i)** Flg22 treatment enhances XBAT35.2-mediated disassociation of VPS28-2 with FLS2. For flg22 treatment, 1uM flg22 was injected into Agro-infiltrated tobacco leaves 30 min before sampling. The experiment was repeated twice with similar results.

The ESCRT machinery plays important roles in trafficking and sorting of internalized PM proteins into MVBs that are usually targeted for vacuolar (in yeast and plants) /lysosomal (in animals) degradation (Paez Valencia et al., 2016). It was previously shown that VPS28-2 and VPS37-1 mediated flg22-induced late endosomal trafficking of FLS2 into the lumen of MVBs (Spallek et al., 2013). Thus, it is conceivable that VPS28-2 and VPS37-1-mediated endosomal trafficking and sorting promotes FLS2 degradation at vacuole. Indeed, when E64D, an inhibitor of lysosomal/vacuolar hydrolases was added, the degradation of FLS2 was significantly inhibited (Fig. 7b, right panel). Importantly, overexpression of VPS28-2 and VPS37-1 also leads to degradation of nonactivated FLS2 proteins and this degradation can also be blocked by E64D treatment (Fig. 7c), further supporting the notion that nonactivated FLS2 proteins also undergo ESCRT-mediated sorting for vacuolar breakdown.

Pathogen growth assays showed that the growth of *Pst* DC3000 in mutant *vps28-2* and *vps37-1.1* was comparable to that in Col-0 whereas the double mutant *vps28-2/vps37-1.1* showed significantly reduced growth of *Pst* DC3000 (Fig. 7d), suggesting negative roles of VPS28-2 and VPS37-1 in plant innate immunity against *Pst* DC3000. This is further substantiated by the observation that overexpression of *VPS28-2* or *VPS37-1* rendered plants more susceptible to *Pst* DC3000 (Fig. 7e). Combining these findings, we conclude that VPS37-1 and VPS28-2 play redundant, negative roles in FLS2-mediated immunity against *Pst* DC3000 by targeting FLS2 for vacuolar degradation.

### XBAT35.2 modulates FLS2 protein level via VPS28-2 and VPS37-1 in flg22- and *Pst* DC3000-triggered plant immunity

We have thus far shown that both XBAT35.2 and VPS28-2 and VPS37-1 are involved in flg22 and *Pst* DC3000-triggered plant immunity by playing opposite roles in regulating FLS2 abundance (Fig. 2a, 2d, 2e, Fig. 4, and Fig. 7a-7e). We have also shown that XBAT35.2 can interact and modify VPS28-2 and VPS37-1 using distinct linkage types of ubiquitin chains (Fig. 5 and Fig. 6a-b), which leads to different outcomes for VPS28-2 and VPS37-1 (Fig. 6c-h). These results suggest that they likely participate in the same physiological pathway with opposite functions. To test this hypothesis, we generate homozygous *35S:VPS28-2-HA*/*xbat35-1* and *35S:VPS37-1-HA*/*xbat35-1* transgenic lines in addition to the *35S:VPS28-2-HA*/Col-0 and *35S:VPS37-1-HA*/Col-0 homozygous line to explore their genetic relationship. We examined the FLS2 protein level in these lines with Col-0 and *xbat35-1* being used as control (Fig. 7f). Without flg22 treatment, overexpression of VPS28-2 or VPS37-1 slightly reduces the FLS2 level in the genetic background of Col-0 and *xbat35-1* compared to that in Col-0, with the reduction in the *xbat35-1* mutant background being slightly more (Fig. 7f upper panel). Activation of immunity by flg22 treatment led to reduction of FLS2 level in all lines tested. However, the reduction was more remarkable in lines where VPS28-2 or VPS37-1 are overexpressed. In particular, the reduction of FLS2 is much more significant in the *xbat35-1* null mutant genetic background (Fig. 7f lower panel). These results together with the findings mentioned above (Fig. 2, Fig.4, Fig. 6, and Fig. 7a-7g) indicated that VPS28-2 and VPS37-1 are epistatic to XBAT35.2 in modulating FLS2 protein level in flg22 and *Pst* DC3000-trigged plant immunity, in which XBAT35.2 regulates FLS2 level via VPS28-2 and VPS37-1.

To further support the conclusion, we next examined the protein stability of VPS37-1-HA under flg22 treatment in transgenic lines where VPS37-1-HA is expressed in Col-0, *xbat35-1,* and *XBAT35.2*-OEX *Arabidopsis* plants. One-hour flg22 treatment led to decline of the VPS37-1-HA protein level in all three genetic backgrounds (Fig. 7g). VPS37-1-HA protein level declined much less in *xbat35-1* than that in the Col-0 whereas to a nearly undetectable level in *XBAT35.2*-OEX lines (Fig. 7g). Degradation assays performed in tobacco leaves also showed that flg22 treatment accelerated XBAT35.2-mediated VPS37-1 degradation (Fig. 7h). Taken these results together with the data shown in Fig. 6, we conclude that flg22 treatment enhances the degradation of VPS37-1 and that heightened XBAT35.2 significantly accelerates the degradation process. Since XBAT35.2-mediated K63-ubiquitination of VPS28-2 does not influence its stability but instead significantly reduces the association of VPS28-2 with FLS2 (Fig. 6d, 6g, and 6h), we analyzed the effect of XBAT35 on the interaction of VPS28-2 with FLS2 in flg22-triggered plant immunity by Co-IP assay. We co-infiltrated the same amount of *A. tumefaciens* harboring constructs that express VPS28-2-FLAG and FLS2CD-HA proteins with increasing amounts of *A. tumefaciens* carrying the XBAT35.2-GFP constructs. As shown in Fig. 7i, the amount of FLS2CD-HA co-immunoprecipitated with VPS28-2-FLAG declined gradually as the amount of XBAT35.2 increased and flg22 treatment significantly boosted this reduction. Thus, these results together corroborate the notion that XBAT35.2 modulates FLS2 protein level via VPS28-2 and VPS37-1 in FLS2-mediated plant immunity.

## DISCUSSION

In the present study, we first identified the tomato XBSL34 (Fti1B) and XBSL35 (Fti1) (Fig. 1, Supplementary Figs. 1 and 2) as interactors of the ubiquitin E2 enzyme Fni3 in efforts to further our understanding of K63-linked ubiquitination in Fen-mediated tomato immunity. The high homology of XBSL35 and XBSL34 to their counterparts in Arabidopsis, XBAT35 and XBAT34, the conservation of XBSL35 and XBAT35 in alternative splicing of their transcript (Supplementary Fig. 3) and their subcellular localization (Fig. 2f and Supplementary Fig. 6a and 6c), and the involvement of XBAT35.2 only in plant immunity against bacterial pathogen (Figs. 2a-e) led us to focus on the role of XBAT35.2 in plant immunity. We revealed that XBAT35.2 interacts with FLS2, BAK1, and BIK1 at the PM *in planta* (Fig. 3a, b) and flg22 treatment enhances the interaction with FLS2 and BAK1 but significantly diminishes the interaction with BIK1 (Fig. 3c-e). These data indicate constitutive association of XBAT35.2 with FLS2, BAK1, and BIK1 at the PM (Fig 2f and Supplementary Fig 6a). A previous study found XBAT35.2 is predominantly localized to Golgi (Liu et al., 2017). Nevertheless, a close look at the confocal microscopic images revealed weaker XBAT35.2-YFP signal around the plasma membrane in that study when XBAT35.2-YFP was expressed in root cells of transgenic Arabidopsis seedlings and tobacco leaf epidermal cells, suggesting localization of XBAT35.2 at PM in that study as well. The negative impact of flg22 treatment on the interaction of XBAT35.2 with the BIK1 is reminiscent of the release of BIK1 from the FLS2/BAK1/BIK1 complex in the early FLS2-mediated immune signaling (Lu et al., 2010). While these data strongly imply engagement of XBAT35.2 in plant immunity initiated by the FLS2/BAK1/BIK1 complex, it is unclear at present the biological significance of the association of XBAT35.2 with FLS2, BAK1, and BIK1 at PM. Since XBAT35.2 is an active ubiquitin E3 ligase, one possible outcome of the interaction is modification of FLS2/BAK1/BIK1 by ubiquitination. However, the FLS2 was not ubiquitinated by XBAT35.2 *in vitro* (Fig. 4g) and overexpression of *XBAT35.2* did not affect the protein level of BAK1 (Fig. 4b). We also did not detect ubiquitination of BIK1 by XBAT35.2 in *in vitro* ubiquitination assay (data not shown). These results, however, cannot completely exclude the possibility that XBAT35.2 is capable of ubiquitinating FLS2 *in planta*. The abundance of XBAT35.2 was suggested to be controlled by autoubiquitination (Liu et al., 2017). Given the findings in this study that XBAT35.2-mediated stabilization of FLS2 is essential for effective FLS2-mediated immunity, another possible outcome for the interaction is phosphorylation of the XBAT35.2 by FLS2, BAK1, and/or BIK1 to block its autoubiquitination and subsequent degradation to promote FLS2-mediated immunity. Further biochemical characterization of the interaction of XBAT35.2 with the FLS2/BAK1/BIK1 complex holds promise for gaining insights into FLS2-mediated early immune signaling at the PM. Through K63 and K48-linked ubiquitination, respectively, XBAT35.2 modifies two different subunits of the ESCRT-I complex, VPS28-2 and VPS37-1 that were shown in this study to mediate the vacuolar degradation of FLS2 (Fig. 7b-c). By thus, XBAT35.2 positively regulates the protein level of FLS2 and consequently, flg22 and bacterial pathogens-triggered plant immunity. Activated FLS2 protein was previously reported to be targeted by PUB12/PUB13 for ubiquitination and subsequent 26S proteasome-dependent degradation to attenuate host immune signaling (Lu et al., 2011). Our findings thus reveal an extra layer of regulation for fine-tuning the protein level of FLS2. Whether the two regulatory circuits for control of FLS2 abundance are connected, and if so, how they are coordinated would be interesting questions to be addressed in next experiments. Given that the ESCRT machinery is involved in an array of cellular and physiological pathways, it is conceivable that the cell must always maintain homeostasis of the machinery for various functions. K63-linked ubiquitination of VPS28-2 reduces its association with FLS2 but not leads to protein turnover, which could facilitate the availability of VPS28-2 for the assembly of the ESCRT-I complex that is crucial for many other cellular functions. In this regard, XBAT35.2-directed distinct ubiquitination of VPS28-2 and VPS37-1 might serve as a mechanism of self-protection for the cell. On the other hand, modification of two subunits of the ESCRT-I complex can help fine-tune the ESCRT-I-mediated sorting of internalized FLS2 for vacuolar breakdown, which in turn is critical for precise regulation of the abundance of FLS2 at the PM. VPS28-2 and VPS37-1 have been implicated in the sorting and delivery of activated FLS2 to the MVB (Spallek et al., 2013). However, XBAT35.2 was not detected at MVB in this study and in a previous study where XBAT35 was shown to target VPS23A, another subunit of the ESCRT-I complex in ABA signaling (Yu et al., 2020). Thus, rather than at MVB, XBAT35.2 likely targets VPS37-1 and VPS28-2 at other location(s), probably before they assemble into the ESCRT-I complex.

Endocytosed non-activated FLS2 proteins can be recycled to the PM through early endosomes or enter Brefeldin A (BFA)-sensitive endosomal pathway (Beck et al., 2012). Our results indicate that elevated XBAT35.2 level increases the FLS2 protein in the absence of flg22 treatment (Fig. 4b), suggesting that part of the internalized non-activated FLS2 proteins may undergo ESCRT-mediated sorting for vacuolar destruction. Consistently, without flg22 treatment, overexpression of VPS28-2 and VPS37-1 reduces the protein level of FLS2 (Fig. 7b, 7c) and the FLS2 protein level is slightly higher in the *vps28-2/vps37-1.1* double mutant compared to that in Col-0 plants (Fig. 7a). Furthermore, in transgenic lines under flg22 treatment, the decrease of FLS2 was significantly less in *vps28-2/vps37-1.1* double mutant (Fig. 7a). And E64D treatment inhibited the vacuolar breakdown of both non-activated and activated FLS2 (Fig. 7b and c). These observations support the notion that both non-activated and activated FLS2 can undergo ESCRT-assisted targeting for vacuole-mediated degradation and flg22-triggered activation of immune signaling enhances ESCRT-I assisted vacuolar degradation of FLS2. Non-activated and activated FLS2 have previously been shown to enter different endosomal pathways (Beck et al., 2012). It would be intriguing to find out in the future how these two forms of FLS2 are targeted by the ESCRT in the endosomal pathway. Regardless, our results presented herein indicate that XBAT35.2 affects FLS2 protein level by modulating subunits of the ESCRT-I complex (Supplementary Fig. 16).

In addition to XBAT35, the XBAT ANK-RING type E3 ligase family also includes four other members, XBAT31-34. Emerging evidence in recent years indicates important regulatory roles for the XBAT proteins in plant growth, development, and responses to various environmental stimuli. XBAT31 regulates *Arabidopsis* hypocotyl growth through mediating the ubiquitination and degradation of the thermosensor ELF3 in response to elevated ambient temperature (Zhang et al., 2021b). XBAT32 was demonstrated to act as a positive regulator of lateral root development through its negative roles in ethylene biosynthesis (Nodzon et al., 2004; Prasad et al., 2010; Prasad and Stone, 2010; Lyzenga et al., 2012). XBAT35 was first characterized in ethylene-mediated control of apical hook curvature in *Arabidopsis* (Carvalho et al., 2012). Subsequent studies have demonstrated the versatility of XBAT35, which is manifested by its broad involvement in plant development, hormone signaling, and biotic and abiotic tolerance (Liu et al., 2017; Li et al., 2020; Yu et al., 2020; Zhang et al., 2021a; Liu et al., 2022). By contrast, no biological functions have so far been assigned to XBAT33 and XBAT34. The five *Arabidopsis* proteins can be grouped into three clades in the phylogenetic tree. Among them, XBAT35 and XBAT34 have the closest evolutionary relationships, with 68% identity and 73% similarity in amino acid sequence (Nodzon et al., 2004). Besides, the two genes also displayed highly similar expression patterns in *Arabidopsis* (Carvalho et al., 2012). We observed that XBAT35.2 and XBAT34 display apparent similarity in their subcellular localization, residing at both the PM and the Golgi (Fig. 2f, and Supplementary Fig. 6a, 6d). These findings suggest XBAT35.2 and XBAT34 may have similar biological functions. However, XBAT35.2 and XBAT34 displayed in this study functional differences in plant immunity against bacterial pathogens. Unlike XBAT35.2, XBAT34 was shown to be not involved in plant defense against bacterial infections, as the *xbat34* mutants showed unaltered immunity to *Pst* DC3000 (Fig. 2a, and Supplementary Fig. 5c). XBAT34 also didn’t interact with FLS2 (Fig. 3a), indicating no direct involvement of XBAT34 in FLS2-mediated immune pathway. However, these results do not preclude XBAT34 from acting in plant innate immunity against other pathogenic microorganisms, such as fungi and viruses.

The ESCRT machinery can be divided into three subcomplexes, ESCRT-I, ESCRT-II, and ESCRT-III and plays a central role in sorting internalized PM proteins for lysosomal/vacuolar destruction (Vietri et al., 2020). In the past four decades, intensive studies have revealed the constituents, structure, assembling mechanisms, and functions of the ESCRT machinery. By contrast, relatively much less is known about how components of the ESCRT complex are modulated to function in various pathways. In mammals, tumor suppressor gene101 (Tsg101) is a key ESCRT-I component that has been shown to be multi-monoubiquitinated or polyubiquitinated (Ferraiuolo et al., 2020). However, evidence that modifications of Tsg101 by ubiquitin directly affect trafficking and sorting of cell surface receptors by the ESCRT-I machinery has not been well established. In plants, VPS23A, the Tsg101 homolog and FREE1 are essential for MVB biogenesis and the endosome-to-vacuole sorting of membrane receptors (Spitzer et al., 2006; Gao et al., 2014). FREE1 was shown to be a crucial regulator in plant response to iron deficiency stress and its homeostasis is mediated by the RING-finger E3 ubiquitin ligases, SINA of *Arabidopsis thaliana* (SINATs) (Xiao et al., 2020). In addition, FREE1 and VPS23A have been known to be involved in the modulation of ABA signaling by transporting and sorting the ABA receptors PYR1/PYL4 for vacuole-mediated turnover (Belda-Palazon et al., 2016; Yu et al., 2016) and to be targeted by SINAT family E3 ubiquitin ligases to promote ABA signaling (Xia et al., 2020). VPS23A was also shown to be targeted by XBAT35 for proteasome-mediated degradation in modulating ABA signaling (Yu et al., 2020). We discover in the present study that the ubiquitin E3 ligase XBAT35.2 modifies VPS37-1 and VPS28-2, two subunits of the ESCRT-I through K48 and K63-linked ubiquitination, respectively, which correspondingly causes degradation of VPS37-1 by the 26S proteasome and diminishment of the interaction between VPS28-2 and FLS2. Interestingly, the rice ANK-RING type ubiquitin E3 ligase XB3 was shown to interact with the rice cell surface immune receptor XA21 and reduced levels of XB3 lead to decreased levels of the XA21 protein and compromised resistance to the avirulent race of *Xanthamonas oryzae* pv *oryzae* (Wang et al., 2006). However, the underlying mechanism for the control of XA21 abundance by XB3 is unknown yet. Based on our findings, we speculate that endocytosis and ensuing trafficking and sorting are also involved in the control of the protein level of XA21 and XB3 targets the ESCRT machinery to modulate XA21 abundance at the PM. More recently, a RING type E3 SGD1 and its homolog that control grain yield in *Setaria italica* and multiple crops was found to ubiquitinate and stabilize the brassinosteroid receptor BRI1 (Tang et al., 2023). The authors speculated that one possible mechanism underlying the stabilization of BRI1 by SGD1 is by targeting the endocytosis pathway. Cell surface receptors play key roles in perceiving and transmitting various types of extracellular signals to internal processes. Our results and the previous reports on VPS23A and FREE1 suggest that modification of ESCRT machinery components by ubiquitin could be a widespread regulatory mechanism for cellular signaling, apparently due to the key roles of the ESCRT machinery in controlling the turnover of membrane proteins, including the numerous cell surface receptors.

## MATERIALS AND METHODS

### Plant Materials

The T-DNA insertion mutant *xbat34* (SALK_132378), *xbat35-1* (SALK_104813), *vps28-2* (SALK_040274c) and *vps37-1.1* (CS804682) are derived from the *Arabidopsis thaliana* ecotype Col-0 whereas the T-DNA insertion mutant *xbat34-#57* (SAIL_395_E02, CS873757) is generated from the *Arabidopsis thaliana* ecotype *Col-3*. All the mutants were ordered from the Arabidopsis Biological Resource Center (ABRC) at the Ohio State University and genotyped with T-DNA and gene-specific primers (Supplementary Table 1). The double mutants (*xbat34*/*xbat35-1* and *vps28-2*/*vps37-1.1*) were generated by crossing corresponding homozygous single mutants. The *35S:VPS28-2-HA* and *35S:VPS37-1-HA*, respectively were introduced into Col-0, *xbat35-1* and *XBAT35.2-FLAG*-overexpressing plants to generate corresponding *VPS28-2* and *VPS37-1*-overexpression lines. Transgenic plants overexpressing XBAT34-FLAG, XBAT35.1-FLAG, XBAT35.2-FLAG were produced by introducing the *35S:XBAT34-FLAG*, *35S:XBAT35.1-FLAG*, and *35S:XBAT35.2-FLAG* construct, respectively into Col-0 plants. *Arabidopsis* seeds were surface-sterilized with 10% (vol/vol) bleach including 0.2% (vol/vol) Triton-100 for 8 min, washed thoroughly in sterile water for 4-6 times before sown in soil or germinated on ½ MS solid medium (pH 5.8-6.0), and grown in the reach-in growth chamber at 22°C and approximately 65% relative humidity under 16 h day/8 h night cycle. *Nicotiana benthamiana* plants were grown on soil in a walk-in growth chamber at 26°C under 16 h day/8 h night long-day cycle.

### DNA Manipulations and Plasmid Constructions

All DNA manipulations were performed according to standard techniques (Green et al., 2012). The constructs for yeast two-hybrid (Y2H) assay using the LexA-based Y2H system (Golemis et al., 2008) were prepared by cloning in frame of the coding sequence of *Fni3*, *Fti1* (*XBSL35*), *Fti1B* (*XBSL34*), *Fti1C*, *Fti1D* and *Fti1E* into the pEG202 bait and pSH18-34 prey vectors, respectively. For protoplast transfection experiments, coding sequences of *XBAT35.2*, *FLS2*, *BAK1* and *BIK1* from their corresponding cDNA were amplified and cloned into the modified pTEX vectors that contain the FLAG or HA epitope tag (Frederick et al., 1998). The constructs for bimolecular fluorescence complementation (BiFC) assay were prepared by PCR amplification of cDNA and cloning the coding sequence of the corresponding genes into the pSPYCE(M) and pSPYNEC 173 vectors (Waadt et al., 2008). To generate constructs for expression and purification of recombinant proteins, coding sequences of the corresponding genes were amplified from cDNA and cloned into the pDEST15 (Thermo Fisher), pDEST17 (Thermo Fisher), and pMAL-c2 (New England Biolabs) to create translational fusion to the GST, 6HIS and MBP epitope tag, respectively. Constructs for overexpressing FLAG-tagged XBAT34, XBAT35.1 and XBAT35.2 in plants were generated by cloning the coding sequences into the pBTEX vector (Frederick et al., 1998). Constructs overexpressing VPS28-2 and VPS37-1 were generated by PCR amplifying and cloning the corresponding coding sequences into the pBTEX1300HA vector in which a HA epitope tag is included to be fused in frame to the protein.

### Immunoblotting and antibody resource

Immunoblotting was performed according to standard techniques (Green et al., 2012) using appropriate antibody. For sample preparation, we weighed and used equal plant tissues for extraction of total proteins and measured the total protein concentration for all samples to ensure equal loading in every experiment. For measuring the relative abundance of the FLS2 protein in immunoblot, the pixel density of the FLS2 band was quantified and normalized to the pixel density of the corresponding Rubisco band using the ImageJ software (Schneider et al., 2012), with the normalized value in untreated Col-0 plants being set as 1. The anti-LexA and anti-FLAG antibodies were ordered from Sigma. The anti-hemagglutinin (HA) (clone 3F10), peroxidase-conjugated monoclonal antibody was purchased from Roche. The anti-ubiquitin (clone P4D1), anti-GST, anti-Myc (clone 9E10), anti-His and anti-GFP antibodies were ordered from Santa Cruz. The anti-FLS2, anti-BAK1, and anti-BIK1 antibodies were purchased from Agrisera. The anti-actin monoclonal antibody (mAbGEa) was ordered from Novus Biologicals. The goat peroxidase-conjugated, anti-rabbit IgG (H + L) secondary antibody and goat peroxidase-conjugated anti-mouse IgG (H + L) secondary antibody were purchased from Bio-Rad.

### Y2H assay

The LexA-based yeast two-hybrid system and procedures used for testing protein-protein interaction were carried out as described previously (Golemis et al., 2008; Mural et al., 2013). Expression of fusion proteins was verified by immunoblotting using either mouse anti-LexA (Sigma) or rat anti-hemagglutinin (HA; 3F10) horseradish peroxidase–conjugated (Roche) monoclonal antibodies.

### Quantitative Real-Time PCR

Seven-day-old *Arabidopsis* seedlings grown on ½ MS plates were immersed in 1 mL sterilized deionized H_2_O in a 24-well plate. After overnight incubation at room temperature, H_2_O was changed into 1μM flg22 solution. Sterilized deionized H_2_O was used as negative control for the flg22 treatment. At the indicated time points, seedlings were sampled and subjected to total RNA extraction using TRI Reagent (Sigma) by following the manufacturer’s protocol. First strand cDNA synthesis was performed using 2ug of total RNA, oligo(dT) 18 primers and the GoScript™ Reverse Transcriptase (Promega). qRT-PCR was performed using gene-specific primers (Supplementary Table 1) and SYBR Green (Life Technologies) on a Bio-Rad iQ5 PCR machine by following the manufacturer’s protocol.

### Pathogen growth assay

The bacterial pathogen *Pst* strain DC3000 cells were cultured on King’s B agar plate with rifampicin (50 mg/L) at 28 °C for 2 days. The bacterial cells were then collected, resuspended in 10 mM MgCl_2_ containing 0.02% Silwet L-77, and diluted to the concentration at OD600 of 0.2. Approximately 4-week-old *Arabidopsis* plants were sprayed with the pathogen inoculum and kept under high humidity for disease development. For measuring bacterial growth, the leaves of inoculated plants were surface sterilized with 70% ethanol for 1 min and rinsed with sterile water for 1 min before three leaf discs were collected using puncher at 2- and 3-day post infection. The leaf discs were thoroughly ground in a 1.5 mL Eppendorf tube and serially diluted and plated on KB plates with rifampicin (25 mg/L) for growing of the pathogen at 28 °C for approximately 2 days, followed by counting the number of colonies and data analysis.

### *Arabidopsis* seedling growth inhibition assay

*Arabidopsis* seeds were surface-sterilized and germinated on ½ MS agar plates. Four-day-old seedlings were transferred into 1 mL ½ MS liquid medium with or without 50 nM flg22 using sterilized 24-well plates. Two seedlings were grown in each well at 24°C and approximately 65% relative humidity under 16 h day/8 h night cycle and the liquid medium was changed every 3 days. The fresh weight of the two seedlings in each well was weighed 10 days after growth in the liquid medium, with a total of 12 seedlings (n=6) being examined for each line.

### Transient protein expression in *Nicotiana benthamiana*

The constructs for transient expression were transformed into *A. tumefaciens* strain GV3101 by electroporation. A 5 mL seed LB culture of the transformed *A. tumefaciens* cells was grown at 28°C overnight with appropriate antibiotics presented. The seed culture was then transferred into suitable amount of LB medium containing appropriate antibiotics, 10 mM MES, and 20 mM acetosyringone the next day for shaker growing at 28°C for 24 h. The cultured bacteria were then collected by centrifugation at 3800g for 15 min. The collected cells were resuspended in infiltration solutions (10 mM MgCl_2_, 10 mM MES, 200 mM acetosyringone) to appropriate OD600 values and kept at room temperature for at least 2 h, followed by Agro-infiltration into 4-week-old tobacco leaves using needleless 1 mL syringes.

### Confocal microscopy

Confocal imaging was carried out using the Nikon A1-NiE upright fluorescence microscope confocal system with the following excitation and emission wavelengths: YFP, 514.5 nm (excitation) and 525 to 555 nm (emission); CFP, 440 nm (excitation) and 480 nm (emission). For subcellular localization analysis, the proteins to be tested and the PM marker (PIP2A-CFP), the ER marker (HDEL-CFP), or the Golgi marker (Man49-CFP) (Nelson et al., 2007) were transiently co-expressed in tobacco leaves by Agro-infiltration. The images were taken 48 hours after infiltration. The subcellular localization of XBAT35.2 was also investigated in Arabidopsis protoplasts by co-transformation of the constructs expressing XBAT35.2-YFP and the PM marker PIP2A-CFP or Golgi marker Man49-CFP using standard protocols. For bimolecular fluorescence complementation (BiFC) assays, the pSPYNE173 and pSPYCE(M) constructs expressing different target proteins were co-introduced into 4-week-old tobacco leaves by Agro-inflitration. The GUS protein was fused with nYFP and cYFP as negative control. The images were taken 48 hours after infiltration.

### Co-immunoprecipitation (Co-IP) Assay

For Co-IP using *Arabidopsis* protoplasts, the protoplasts were prepared and transfected as described previously (Yoo et al., 2007). In brief, 500 μL protoplasts were transfected with 40-50 μg of indicated plasmids, incubated overnight at room temperature, and then treated with H_2_O or 1μM flg22 for 10 min. For Co-IP using tobacco leaves, different combinations of *A. tumefaciens* carrying the constructs of interest were co-infiltrated into tobacco leaves. GFP was used as internal expression control. Total proteins were then extracted 2 days after infiltration using Co-IP extraction buffer containing 50 mM Tris-HCl, pH 7.5, 150 mM NaCl, 10% glycerol, 10 mM DTT, 1 mM NaF, 1 mM Na_2_MoO_4_·2H_2_O, 1% (v/v) Phosphatase Inhibitor Cocktails (Sigma-Aldrich), 1% (v/v) Protease Inhibitor Cocktail (P9599, Sigma-Aldrich), 100 μM phenylmethylsulphonyl fluoride and 0.5% (v/v) Triton-100. Supernatants were incubated for 3-4 h at 4 °C with 15-20 μL of ANTI-FLAG® M2 Affinity Gel (Sigma) and washed 3-4 times with Co-IP wash buffer containing 0.1% Triton-100 but without Phosphatase Inhibitor Cocktails and Protease Inhibitor Cocktail. The immunoprecipitated protein sample was incubated in 1xSDS loading buffer at 95 °C for 5 min, centrifuged for 1 min at 14, 000g, and then the supernatants were analyzed by immunoblotting with an anti-HA or anti-FLAG antibody.

### Protein degradation Assay

For examining the possible degradation of VPS37-1 and VPS28-2 by XBAT35.2 *in planta*, the experiments were performed as previously described with some modifications (Yu et al., 2020). In brief, different combinations of *A. tumefaciens* carrying constructs of interest were co-infiltrated into tobacco leaves. GFP was used as internal expression control. Leaf samples were collected 2 days after the infiltration. Total proteins were then extracted from the leaves using protein extraction buffer employed for the Co-IP assays and subjected to immunoblotting using anti-HA, anti-FLAG, and anti-GFP antibodies, respectively. Coomassie Brilliant Blue (CBB) staining for the Rubisco proteins was used as loading control.

### Recombinant protein expression and purification

All GST-tagged (pDEST15), His-tagged (pDEST17), and MBP-tagged (pMAL-C2) fusion proteins were expressed using the BL21 *Escherichia coli* strain and the affinity purification was performed using standard protocols. The protein concentration was determined using Bio-Rad protein assay dye reagent concentrate (Bio-Rad).

### *In vitro* and *in vivo* ubiquitination assays

The *in vitro* ubiquitination assay was performed as described previously (Mural et al., 2013). In brief, the reaction was performed in 30 μL ubiquitination buffer (50 mM Tris-HCl, pH 7.5, 5 mM ATP, 5 mM MgCl_2_, 2 mM DTT, 3 mM Creatine phosphate, 5 μg/ml Creatine phosphokinase) containing 10 μg of ubiquitin, 40 ng of E1, 200 ng of E2, 3 μg of E3 and 200 ng of substrates at 30 °C for 3 h. The reactions were stopped by adding 10 μl 4× SDS loading buffer and heating at 95 °C for 5 min. 5-10 μL of reaction sample were used for immunoblotting analysis. For detecting the ubiquitination of VPS28-2 and VPS37-1 by XBAT35.2 *in planta*, the seedlings of Arabidopsis transgenic lines where VPS28-2 or VPS37-1-HA expressed in the genetic background of Col-0, *xbat35-1*, and *35S:XBAT35.2-FLAG*, respectively were grown on ½ MS medium plus agar plates for 10 days under long-day condition, then transferred to a flask with liquid ½ MS medium containing 75 μM MG132 and shaken overnight. Total proteins were extracted from the MG132-treated seedlings using the extraction buffer as previously described with minor modification by adding 75 μM MG132 (Aguilar-Hernández et al., 2017), followed by immunoblotting analysis using anti-HA antibodies.

### Data statistical analysis

Statistical significance among treatments was determined by one-way ANOVA (Tukey HSD), and asterisks indicate the statistical significance: *: p< 0.05, **: p< 0.01.

## ACKNOWLEDGMENTS

We thank Christine Elowsky, Terri Fangman, and You “Joe” Zhou (Biotechnology Center, University of Nebraska-Lincoln) for helping with the confocal microscope. This research was supported, in part, by start-up funds from the University of Nebraska-Lincoln and grant from the United States Department of Agriculture (2012-67014-19449) and National Science Foundation (IOS-1645659) to LZ.

---

**Supplementary Figure 1.**
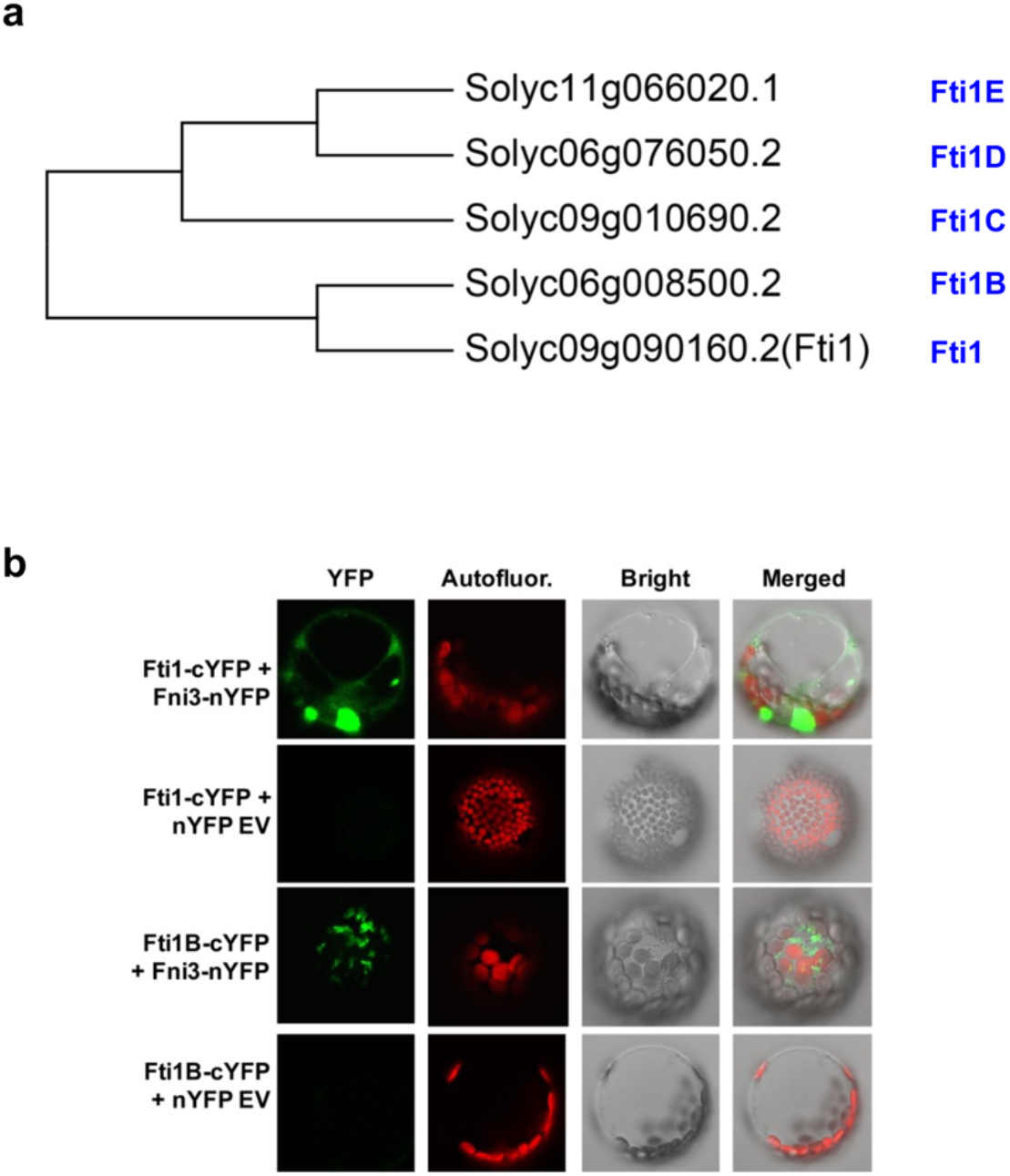
Fti1 (XBSL35) and Fti1B (XBSL34) interact with Fni3 *in vivo*. (**a**) Phylogenetic tree of Fti1 family members. The tree was constructed by neighbor-joining method using the MEGA11 software with 500 bootstrap replicates. (**b**) The interaction of Fti1 and Fti1B with Fni3 was confirmed by BiFC assay using tomato leaf protoplasts. The protoplasts in which C-terminal YFP-fused Fti1 (Fti1-cYFP) or Fti1B (Fti1B-cYFP) and N-terminal YFP empty vector (nYFP-EV) co-expressed were used as negative control.

**Supplementary Figure 2.**
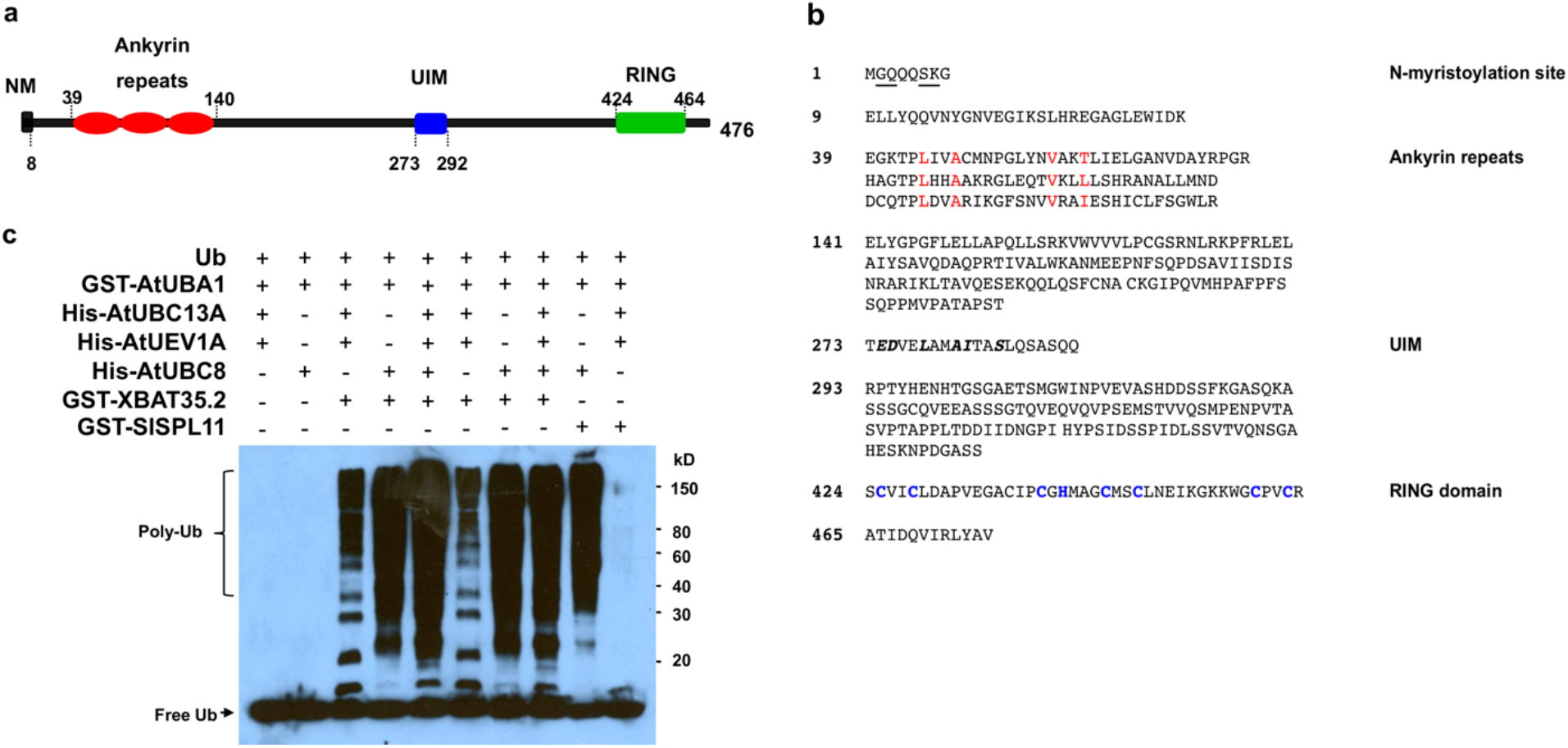
Fti1 (XBSL35) and Fti1B (XBSL34) are ANK-RING type proteins. (**a**) Schematic representation of domain organization for the Fti1 protein. NM, putative N-myristoylation site; UIM, ubiquitin-interacting motif. (**b**) Protein sequence of Fti1 with key motifs and domains being indicated. The conserved amino acid residues for the N-myristoylation site, the Ankyrin repeats, the UIM motif, and the signature amino acids C3HC4 for the RING domain are underlined, in red color, in italic, and in blue color, respectively. (**c**) XBAT35.2 work with E2s UBC8 and UBC13A to catalyze K48 and K63-linked autoubiquitination, respectively. SlSPL11, the tomato homolog to the rice SPL11 ubiquitin E3 ligase was used as a positive control. The numbers on the right show the molecular mass of marker proteins in kilodaltons. Poly-Ub: polyubiquitin chains. This experiment was repeated twice with similar results.

**Supplementary Figure 3.**
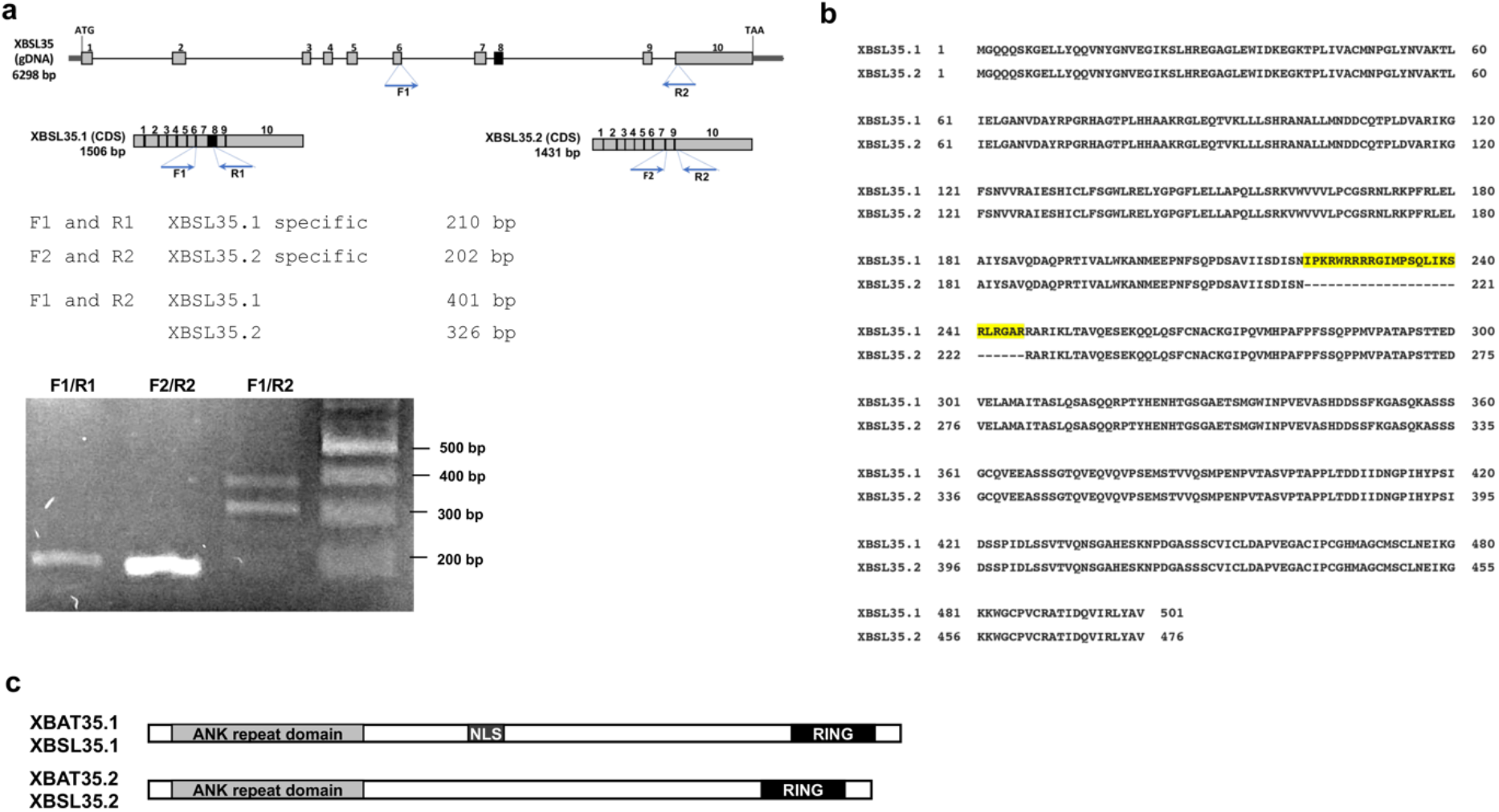
The *Arabidopsis XBAT35* and tomato *XBSL35* genes are conserved in alternative splicing that each produces two isoforms. (**a**) Upper panel: Schematic illustration of the exons in the genomic DNA and CDS of two isoforms for the tomato *XBSL35* gene. Lower panel: Reverse transcriptase (RT)-PCR amplification of the region that spans the exon 8 encoding a nuclear localization signal (NLS) sequence using the two transcript isoforms of *XBSL35* as template. (**b**) Alignment of the protein sequence for the two isoforms of the XBSL35 protein, XBSL35.1 and XBSL35.2. The sequence alignment was performed using Clustal X. (**c**) The two protein isoforms of XBAT35 and XBSL35 have conserved domain organization. The difference between XBSL35.1 and XBSL35.2, as well as between XBAT35.1 and XBAT35.2 is a NLS sequence.

**Supplementary Figure 4.**
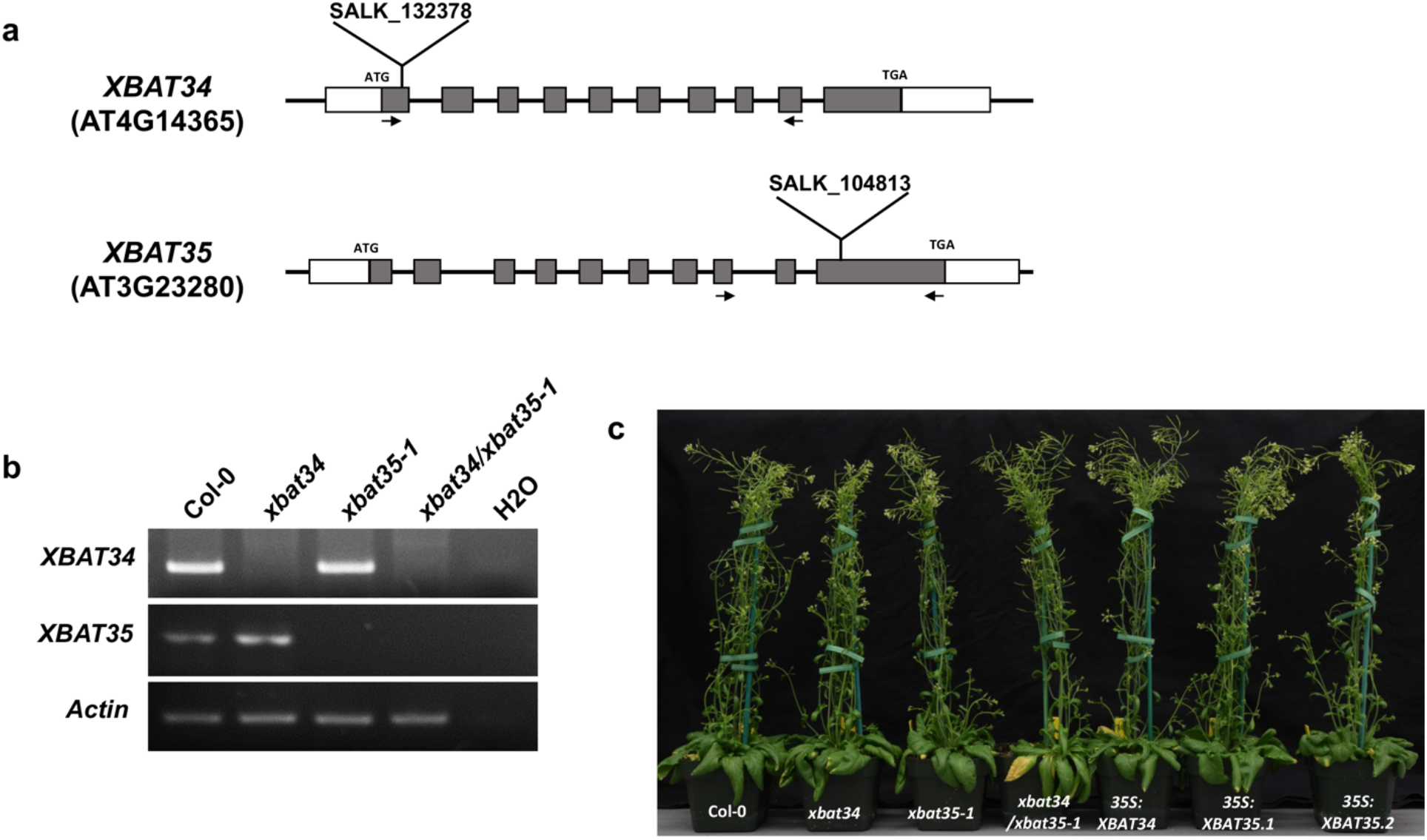
Identification of the *xbat34* and *xbat35-1* T-DNA insertion mutants. (**a**) Schematic representation of *XBAT34* and *XBAT35* genomic structure with the positions of the T-DNA insertion site being shown. Gray-filled boxes: exons; White boxes: 5’ and 3’ UTR. Filled arrows denote the position of primers used for examining the expression of corresponding gene in (b). (**b**) Examination of the expression of *XBAT34* and *XBAT35* in the indicated T-DNA insertion mutant lines by RT-PCR. Gene-specific primers were used to amplify the transcripts of each gene from the cDNA samples. The *Actin* gene was used as control. This experiment was repeated twice with similar results. (**c**) Growth phenotype of Col-0, *xbat34*, *xbat35-1* and *xbat34/xbat35-1* mutants and *XBAT34*, *XBAT35.1* and *XBAT35.2*-overexpressing *Arabidopsis* plants. Photo was taken for approximately 5-week-old plants.

**Supplementary Figure 5.**
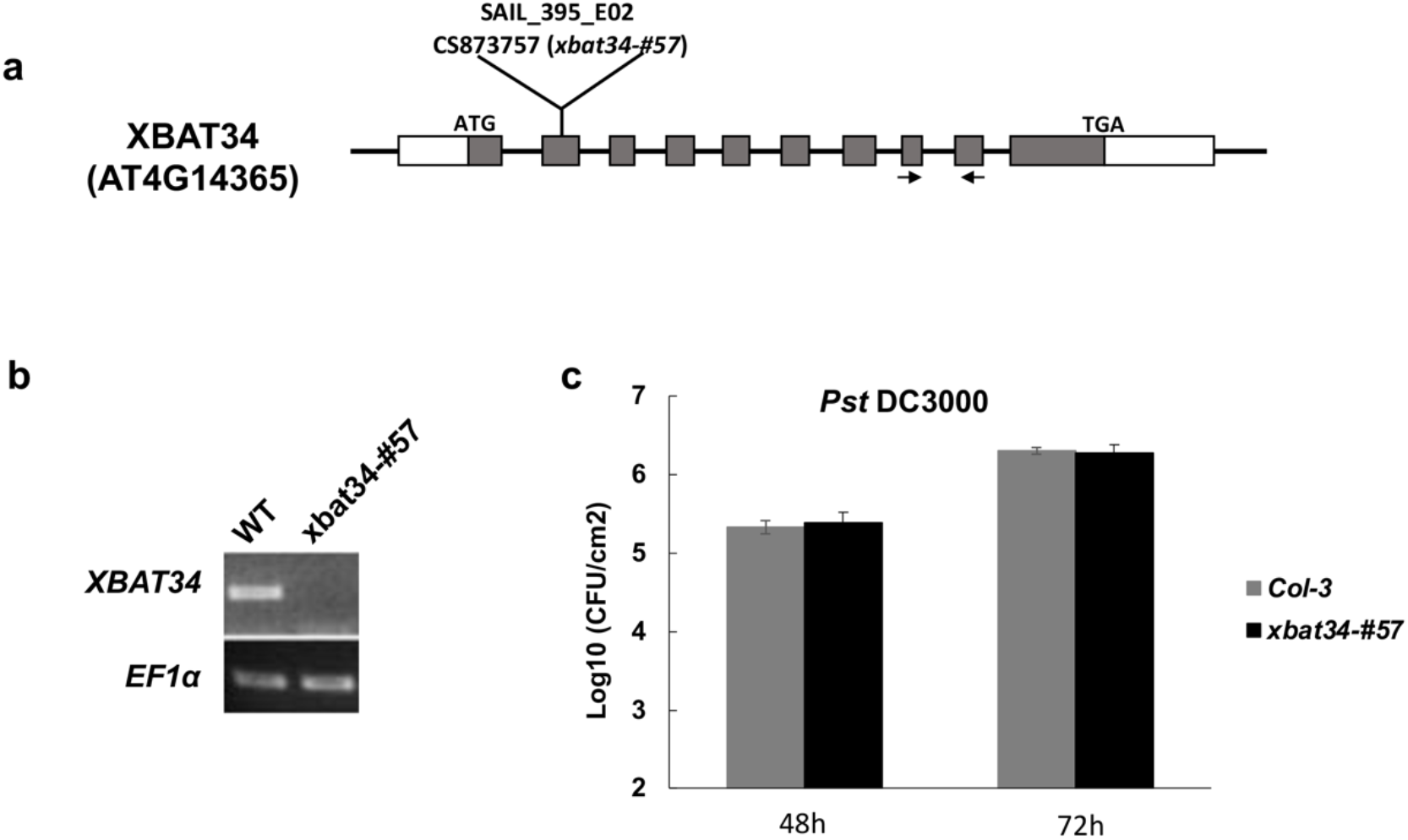
A T-DNA insertion mutant of *XBAT34* in Col-3 background, *xbat34-#57* displays comparable immunity as the wild-type plant to *Pst* DC3000. (**a**) Schematic representation of *XBAT34* genomic structures with the position of the T-DNA insertion in *xbat34-#57* being shown. Gray-filled boxes: exons; White boxes: 5’ and 3’ UTR. Filled arrows denote the position of primers used for examining the expression of *XBAT34* in (b). (**b**) Examination of the *XBAT34* gene expression in Col-3 and the *xbat34-#57* mutant line by RT-PCR. Gene-specific primers were used to amplify the transcripts of each gene from the cDNA samples. The *EF1a* gene was used as control. (**c**) Growth of *Pst* DC3000 on Col-3 and *xbat34-#57* plant leaves. Four-week-old plants were inoculated with *Pst* DC3000 (OD600=0.2), and the bacterial growth was determined at 48 and 72 hpi. Error bars represent the SD (n=9). This experiment was repeated twice with similar results.

**Supplementary Figure 6.**
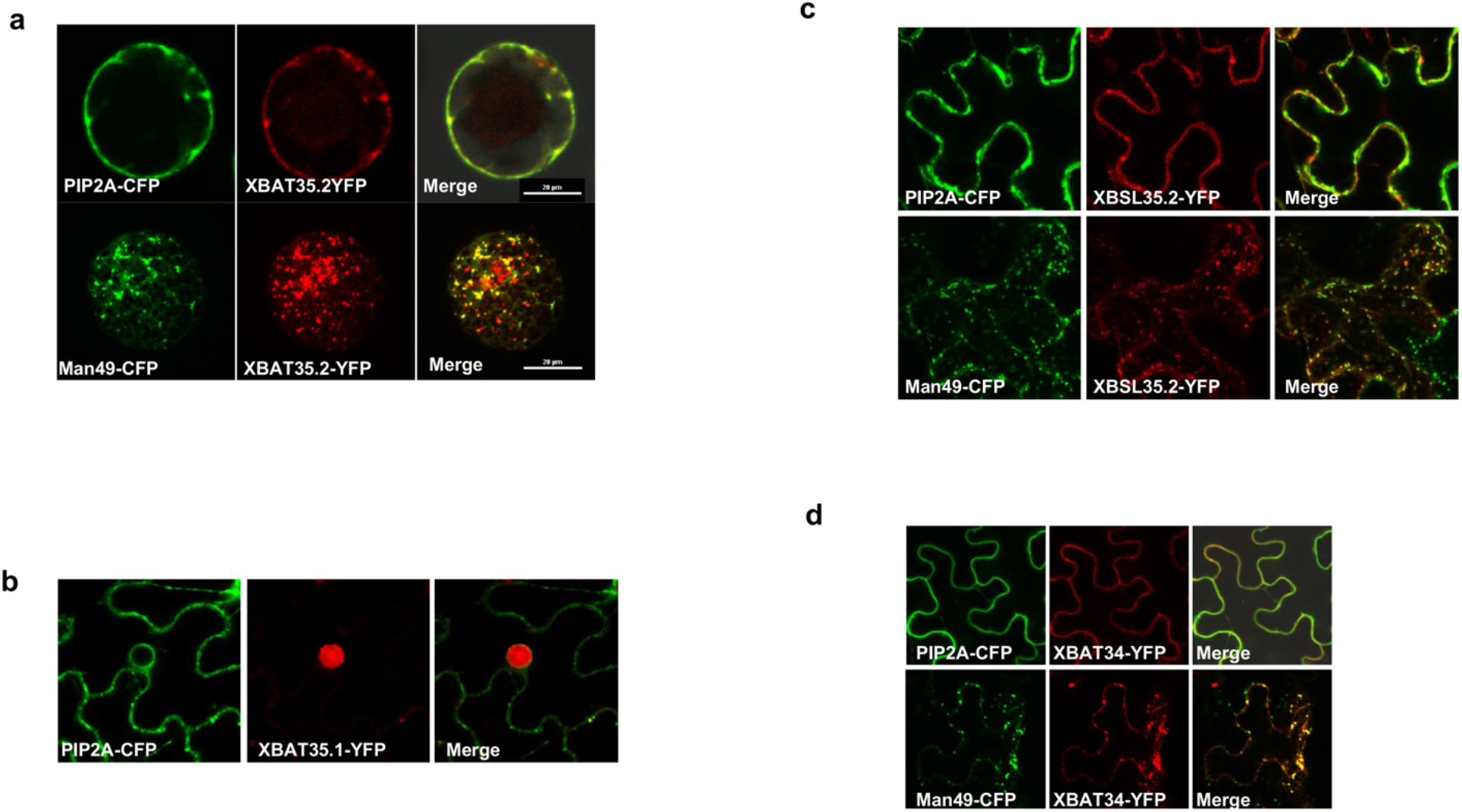
Subcellular localization of XBAT34, XBAT35.1 and XBSL35.2. (**a**) Subcellular localization of XBAT35.2 in *Arabidopsis* leaf-derived protoplasts. The YFP signal is pseudo-colored in red, and CFP signals are shown in green. PIP2A:CFP and Man49:CFP are PM and Golgi-specific marker, respectively (Nelson et al., 2007). White bar, 20 micrometers (μm). This experiment was repeated twice with similar results. (**b**) XBAT35.1-YFP is located in the nucleus and doesn’t colocalize with the PM marker (PIP2A-CFP) in tobacco leaf epidermal cells. YFP signals are pseudo-colored in red. CFP signals are shown in green. (**c**) XBSL35.2-YFP colocalizes with the PM marker (PIP2A-CFP) and Golgi marker (Man49-CFP) in tobacco leaf epidermal cells. The YFP signals are pseudo-colored in red, and CFP signals are shown in green. (**d**) XBAT34-YFP colocalizes with the PM marker (PIP2A-CFP) and Golgi marker (Man49-CFP) in transiently co-transformed tobacco leaf epidermal cells. Experiments in figures (a), (b), and (c) were repeated at least twice with similar results.

**Supplementary Figure 7.**
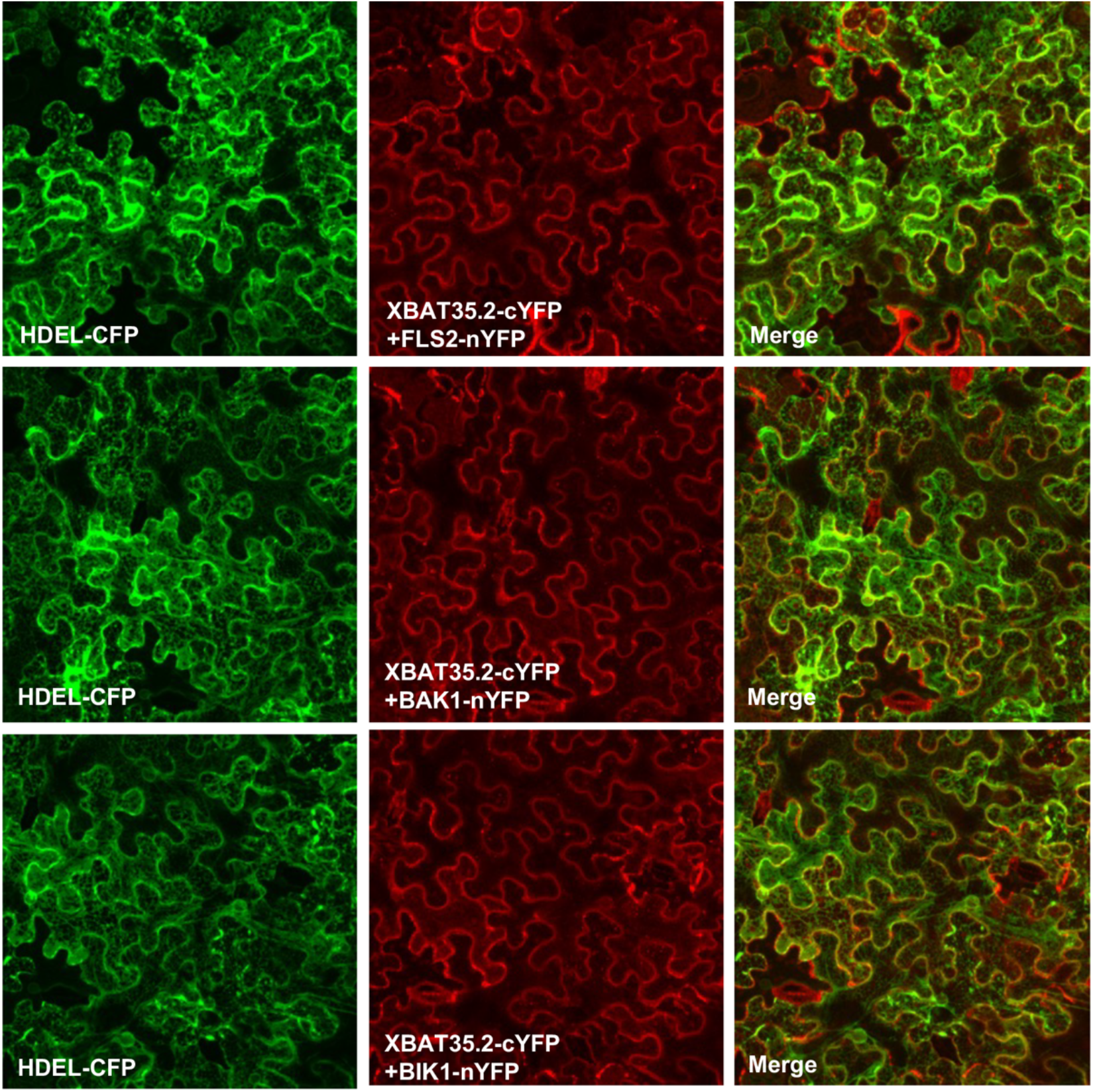
Interaction of XBAT35.2 with FLS2, BAK1, and BIK1 does not occur at the endoplasmic reticulum (ER). An ER-marker, HEDL-CFP (Nelson et al., 2007) was added to the BiFC assay for the interaction of XBAT35.2 with FLS2, BAK1, and BIK1 in tobacco leaf. The CFP signal (pseudo-colored in green) and YFP signal (pseudo-colored in red) were detected in epidermal cells. The YFP signal was detected 48 h after infiltration. This experiment was repeated twice with similar results.

**Supplementary Figure 8.**
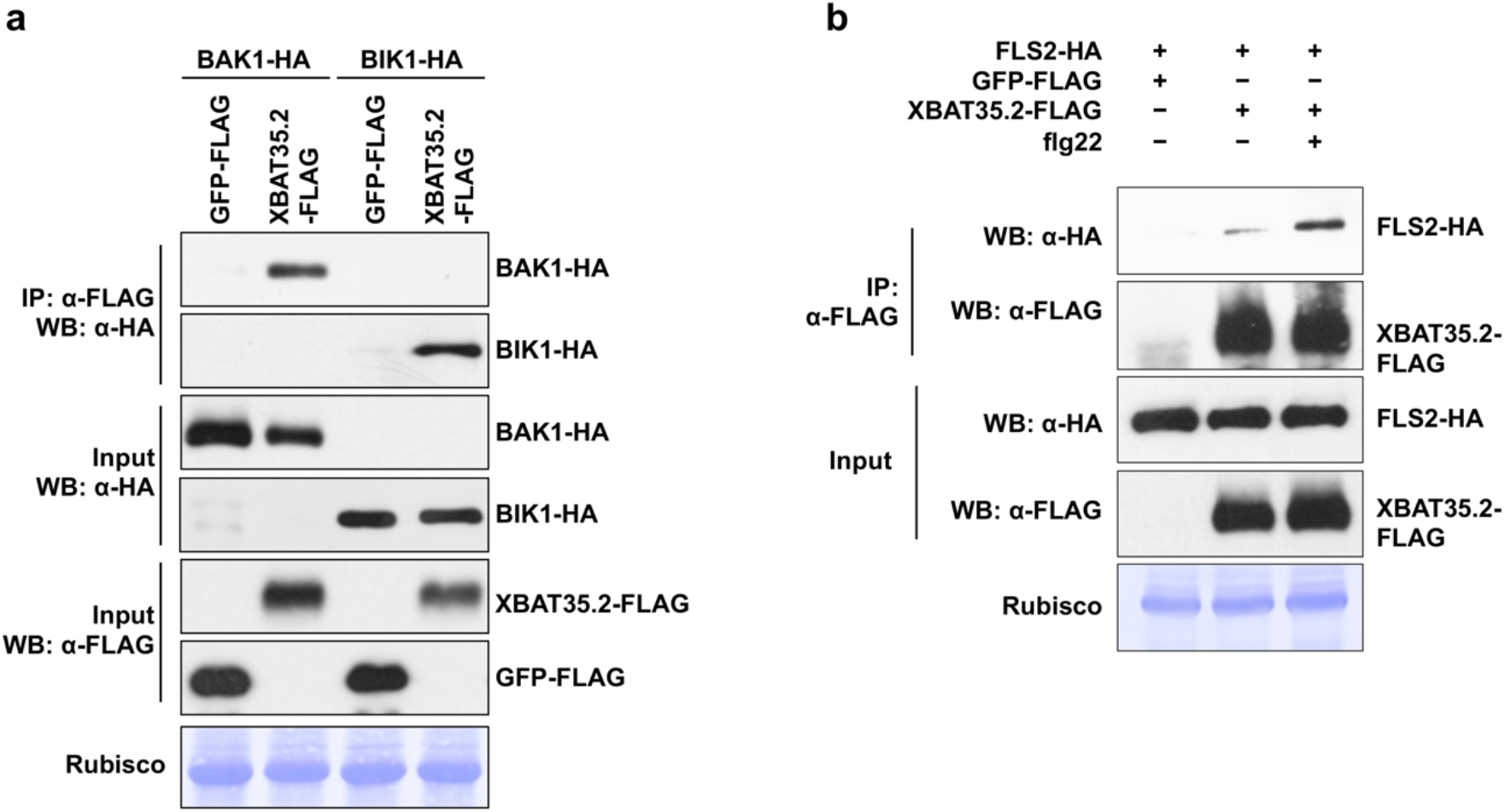
Interaction of XBAT35.2 with FLS2, BAK1, BIK1 in tobacco leaves. (**a**) Co-IP assay was performed using XBAT35.2-FLAG, BAK1-HA and BIK1-HA transiently co-expressed in tobacco leaves. Stained rubisco bands denote equal loading. GFP-FLAG was used as negative control. This experiment was repeated twice with similar results. (**b**) Co-IP assay was performed using XBAT35.2-FLAG and FLS2-HA transiently co-expressed in tobacco leaves. GFP-FLAG was used as negative control. Stained rubisco bands denote equal loading. For flg22 treatment, tobacco leaves were infiltrated with 1μM flg22 for 30 min before sampling.

**Supplementary Figure 9.**
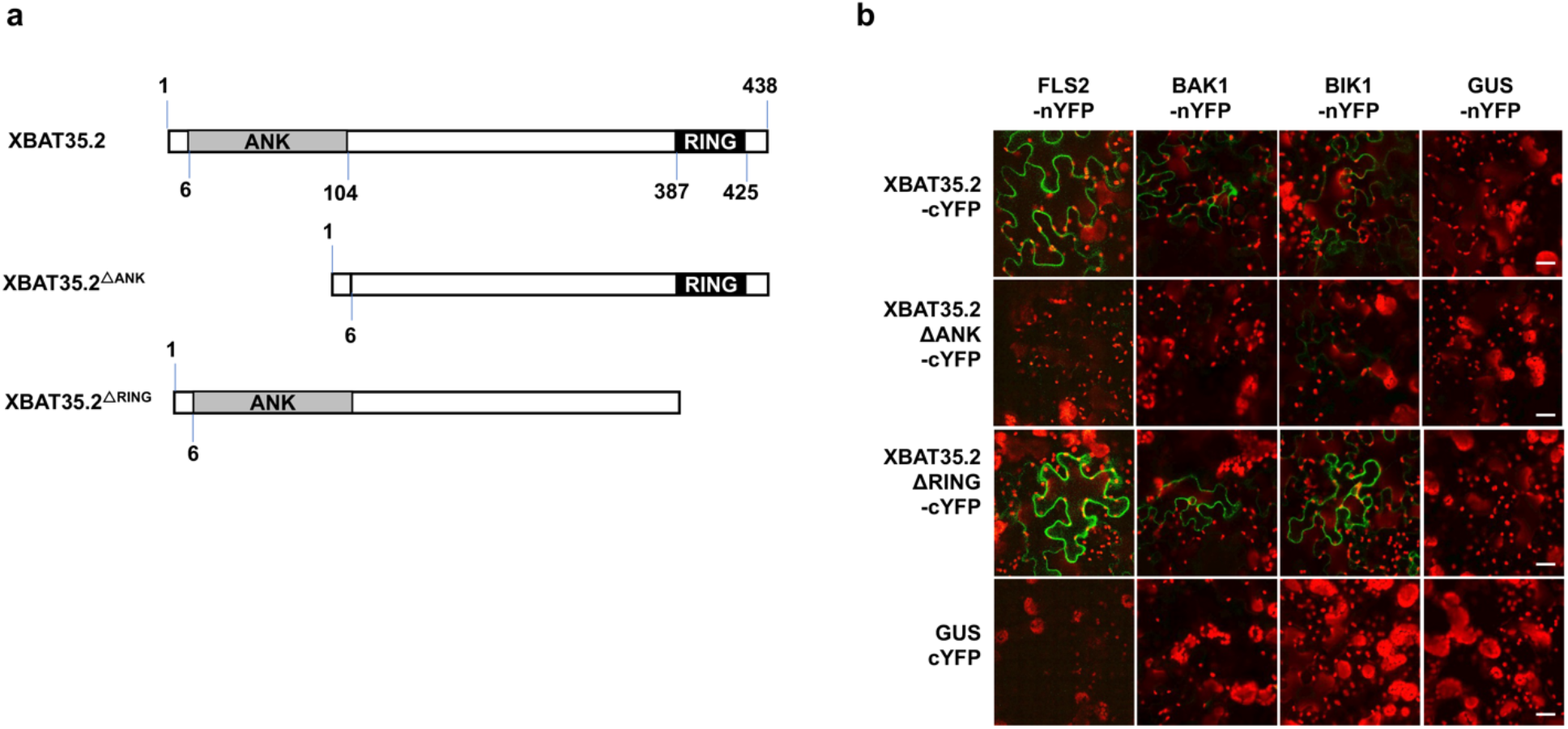
The Ankyrin repeats (ANK) domain of XBAT35.2 is essential for interaction with FLS2, BAK1, and BIK1. (**a**) Schematic representation of the wild type and truncated versions of XBAT35.2. The numbers denote position of the start and end amino acid of the corresponding domain. (**b**) BiFC assay of the interaction of wild type and truncated versions of XBAT35.2 with FLS2, BAK1, and BIK1 in tobacco leaf epidermal cells. YFP signal (pseudo-colored in green) indicates the interaction of corresponding co-expressed proteins. The YFP signal was detected 48h after Agro-infiltration. GUS was used as negative control. White bar, 20 micrometers (μm). This experiment was repeated twice with similar results.

**Supplemental Figure 10.**
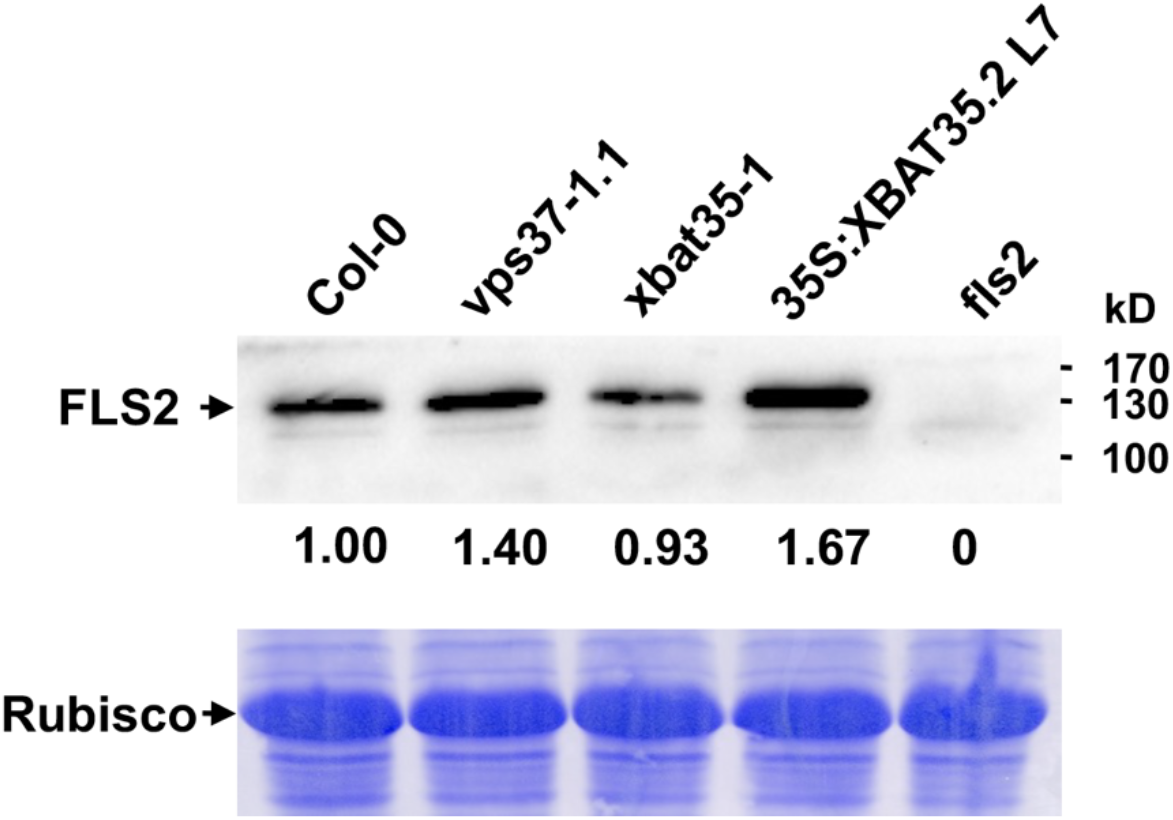
The FLS2 level in four-week-old Arabidopsis plants grown in soil. The protein level of FLS2 in Col-0, *vps37-1.1*, *xbat35-1* and XBAT35.2-overexpressing (line #7), and the fls2 null mutant Arabidopsis plants was examined using the anti-FLS2 antibody ordered from Agrisera. The plants were grown in soil for 4 weeks. Two 0.8 mm-diameter leaf discs were then sampled from the fully expanded leaves, followed by total proteins extraction and immunoblotting analysis. The numbers below the bands denote the relative abundance of the FLS2 protein by quantifying the pixel density of the FLS2 band and normalizing to the pixel density of the corresponding Rubisco band using the ImageJ software (Schneider et al., 2012), with the normalized value in untreated Col-0 plants being set as 1. The specificity of the anti-FLS2 antibody is manifested by the absence of the FLS2 immunoblot signal in the fls2 mutant.

**Supplementary Figure 11.**
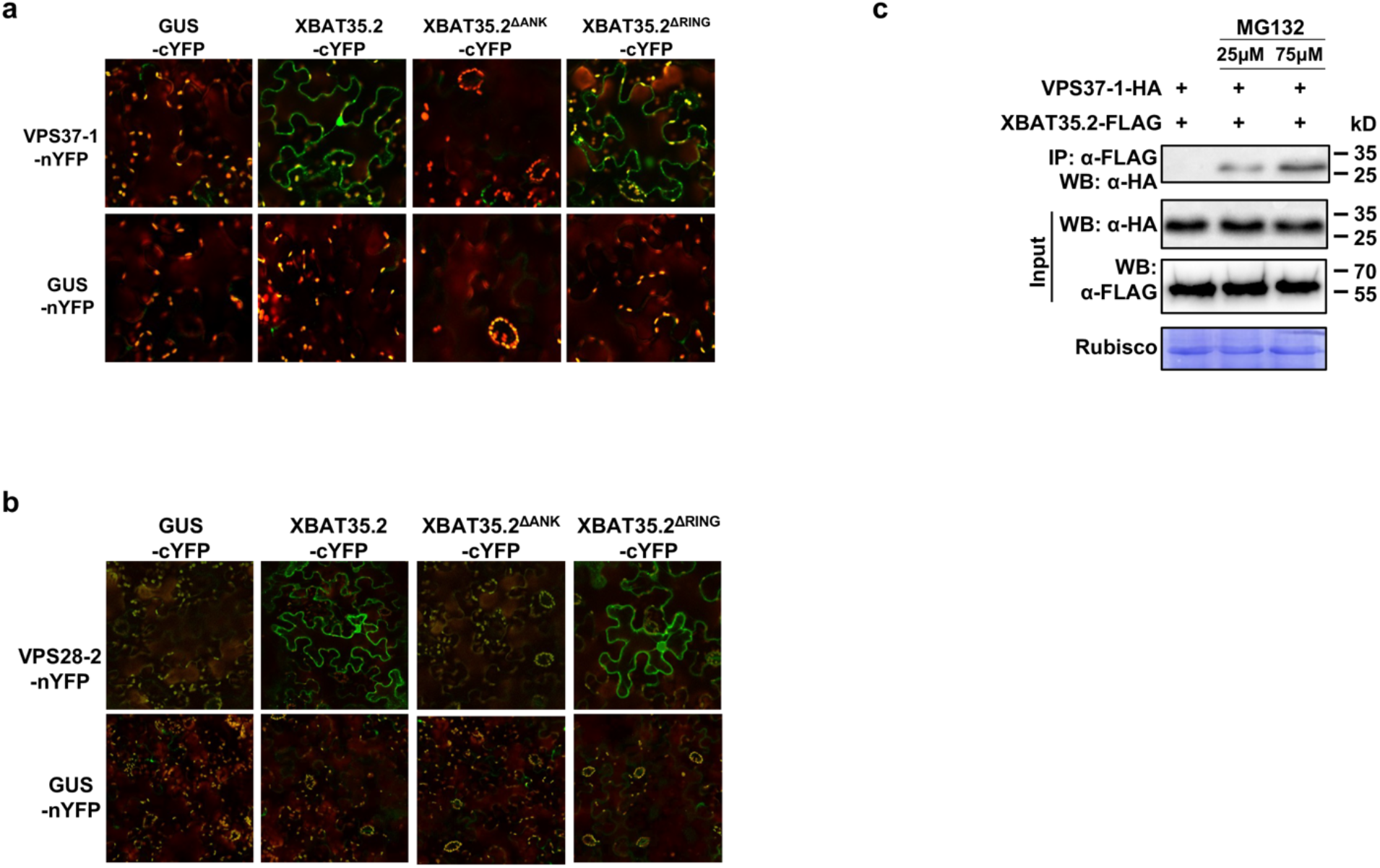
Interaction of XBAT35.2 with VPS28-2 and VPS37-1 *in vivo*. (**a**) and (**b**) BiFC analysis of the interaction of nYFP-fused VPS37-1 (a) and VPS28-2 (b) with cYFP-fused XBAT35.2 and its truncation mutants (XBAT35.2^ΔANK^ and XBAT35.2^ΔRING^) in tobacco leaf epidermal cells. YFP signal (pseudo-colored in green) indicates interaction between the two proteins tested. The tobacco leaves were collected and examined on the third day after Agro-infiltration. GUS and FLS2 were used as negative and positive control, respectively. The experiments were repeated twice with similar results. (**c**) MG132 treatment promotes the coprecipitation of VPS37-1 with XBAT35.2. *A. tumefaciens* cells harboring the constructs that express VPS37-1-HA and XBAT35.2-FLAG were co-infiltrated into tobacco leaves. Total protein was extracted from leaves on the third day after Agro-infiltration. Co-immunoprecipitation was carried out with anti-FLAG antibody, and the immuno-precipitated samples were analyzed by immunoblotting using anti-HA antibody.

**Supplementary Figure 12.**
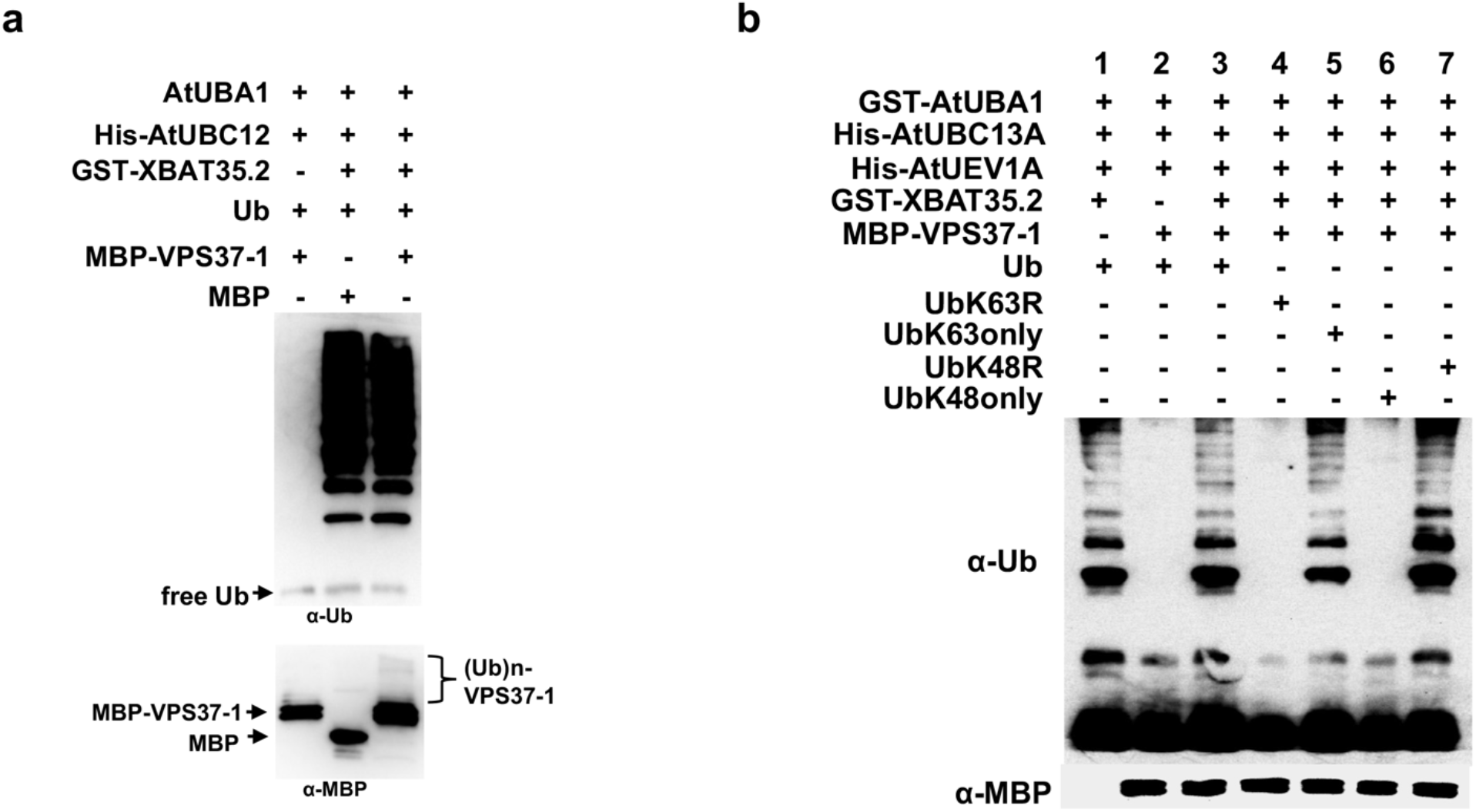
XBAT35.2 does not ubiquitinate VPS37-1 with K63-linked ubiquitin chains and targets VPS37-1 for degradation. (**a**) XBAT35.2 ubiquitinates VPS37-1 *in vitro*. Upper panel: XBAT35.2 autoubiquitination detected by anti-ubiquitin antibody denotes E3 ligase activity of XBAT35.2. Lower panel: The high molecular weight smear marks ubiquitination of VPS37-1 [(Ub)n-VPS37-1]. MBP was used as negative control. This experiment was repeated three times with similar results. (**b**) *In vitro* ubiquitination of MBP-VPS37-1 by XBAT35.2 was assayed in the presence of an *Arabidopsis* E1 (AtUBA1), E2 (AtUBC13a), E2 variant AtUEV1a. Upper panel: XBAT35.2 autoubiquitination detected by anti-ubiquitin antibody. Lower panel: anti-MBP antibody was used to detect the ubiquitination of MBP-VPS37-1.

**Supplementary Figure 13.**
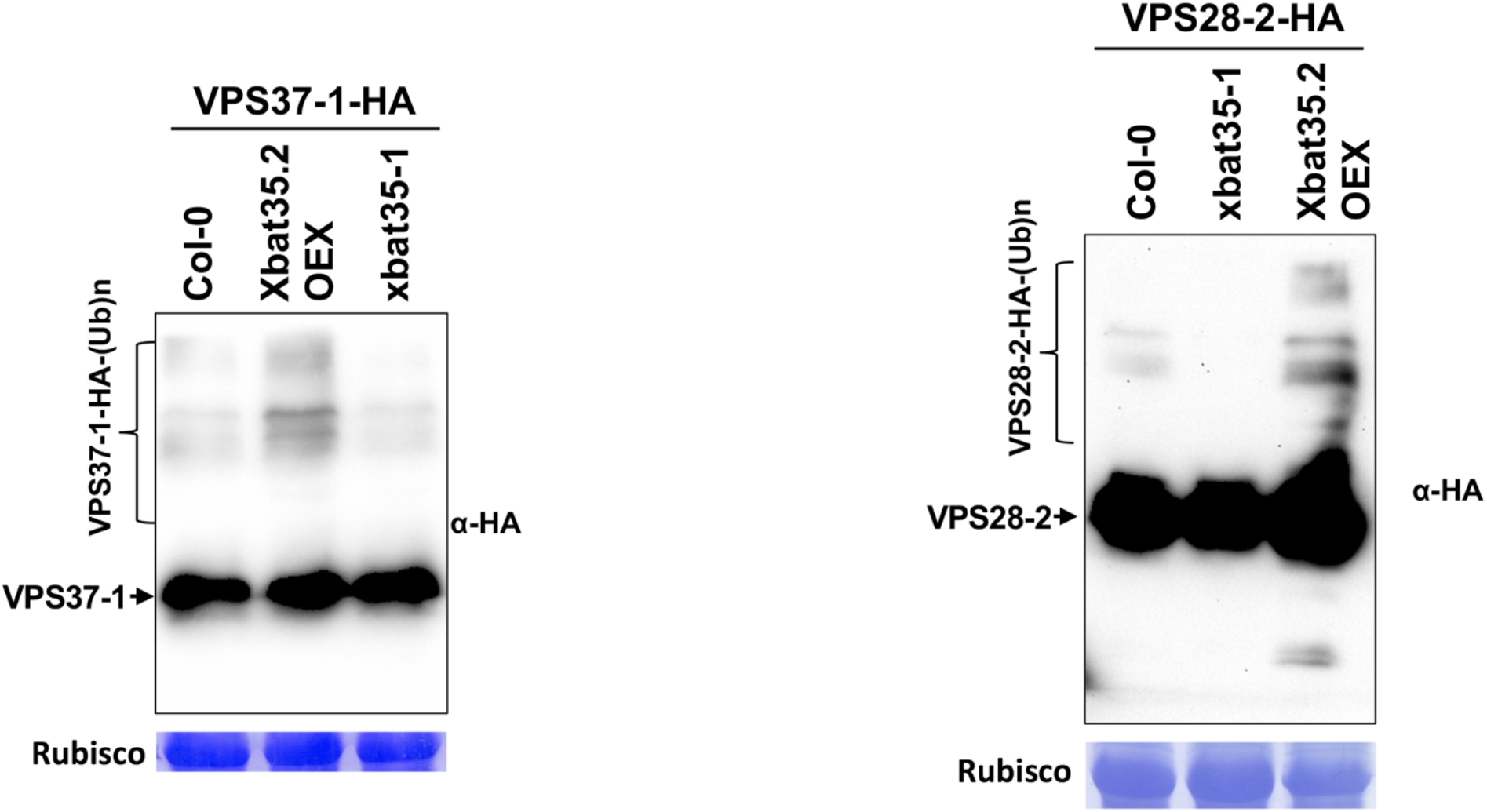
XBAT35.2 directs ubiquitination of VPS28-2 and VPS37-1 *in vivo.* The ubiquitination of VPS37-1 (A) and VPS28-2 (B) *in vivo* were detected in seedlings of Arabidopsis transgenic lines where VPS28-2 or VPS37-1-HA expressed in the genetic background of Col-0, *xbat35-1*, and *35S:XBAT35.2-FLAG*, respectively. The seedlings were grown on ½ MS medium plus agar plates for 10 days under long day condition, then transferred to a flask with liquid ½ MS medium containing 75 μM MG132 and shaken overnight before total proteins extraction and immunoblotting.

**Supplementary Figure 14.**
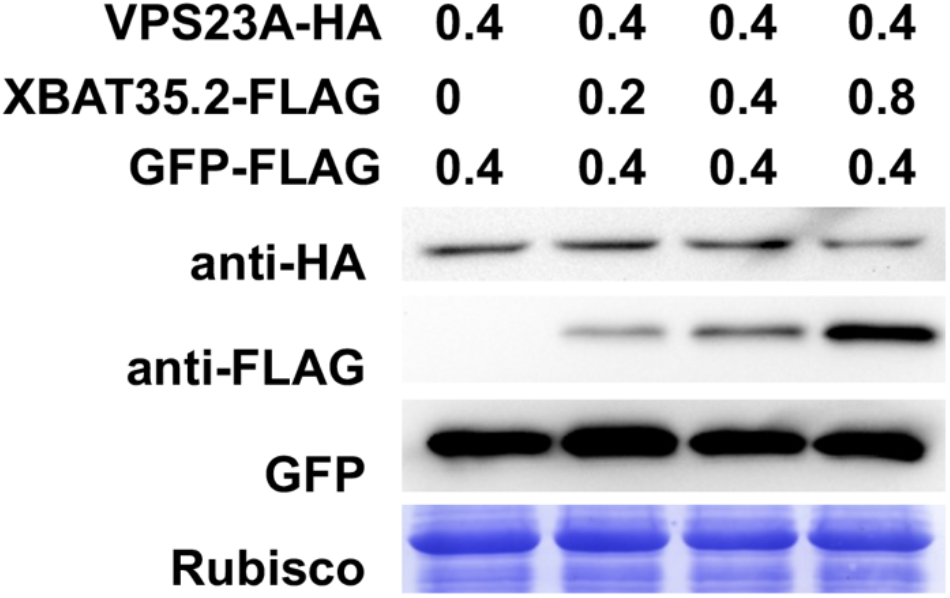
XBAT35.2 promotes degradation of VPS23A. *A. tumefaciens* cells carrying the *35S:VPS23A-HA* construct were co-infiltrated into tobacco leaves with increasing amount of *A. tumefaciens* cells that express XBAT35.2-FLAG. GFP-FLAG was used as internal expression control. Samples were collected 48 h after infiltration, followed by immunoblotting analysis using anti-HA antibodies. This experiment was repeated twice with similar results.

**Supplementary Figure 15.**
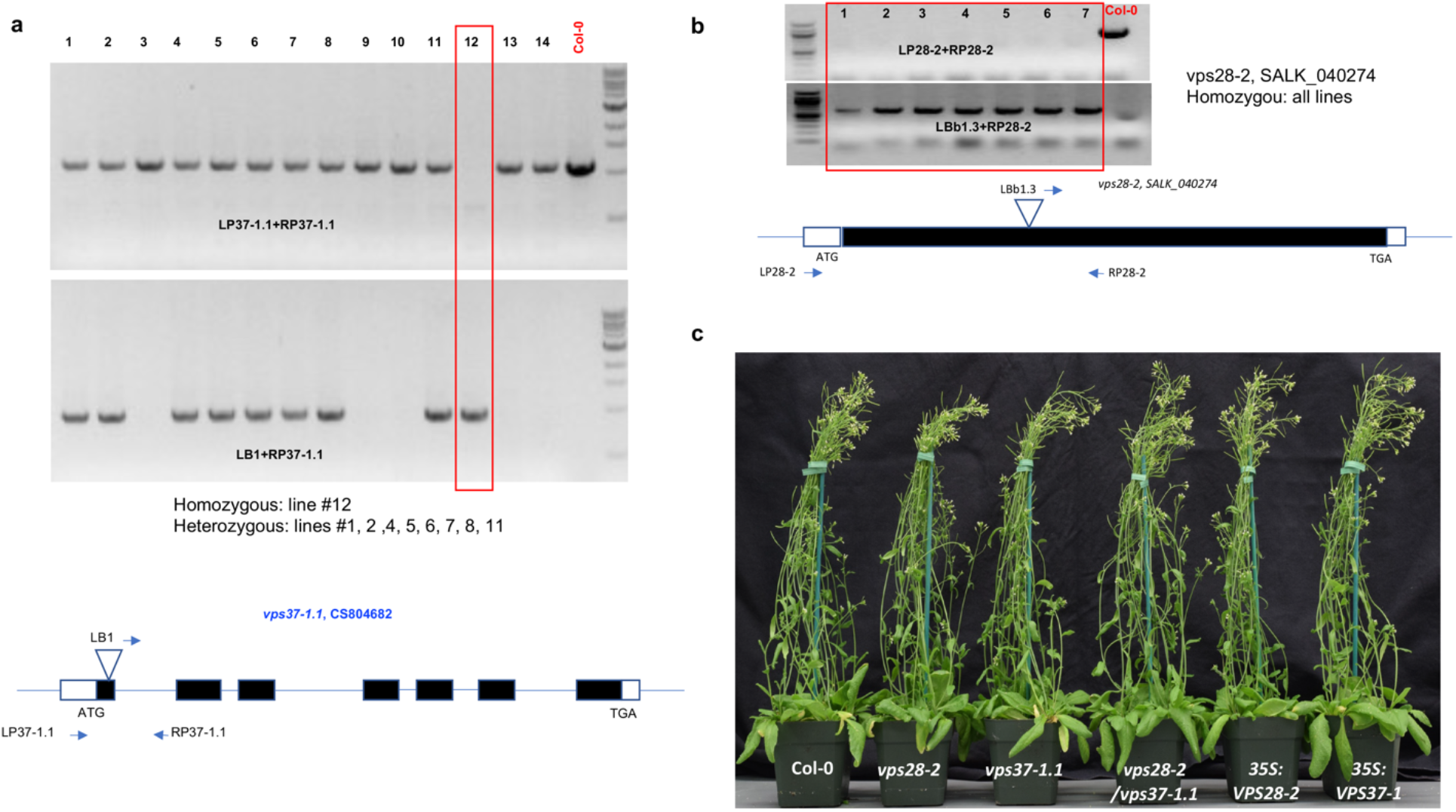
Identification of the *vps28-2* and *vps37-1.1* T-DNA insertion mutants. (**a**) Identification of the *vps37-1.1* T-DNA insertion mutant. Upper panel: genotyping the T-DNA insertion using primers indicated in the lower panel. Lower panel: schematic representation of the genomic structure of *VPS37-1* with the position of the T-DNA insertion being shown. The gene-specific primers and a primer at the right border of the T-DNA for genotyping are indicated as filled arrows. Filled boxes: exons; open boxes: 5’ and 3’ UTR region. (**b**) Identification of the *vps28-2* T-DNA insertion mutant. Upper panel: genotyping the T-DNA insertion using primers indicated in the lower panel. Lower panel: schematic representation of the genomic structure of *VPS28-2* with the position of the T-DNA insertion being shown. The gene-specific primers and a primer at the right border of the T-DNA for genotyping are indicated as filled arrows. Filled boxes: exon; open boxes: 5’ and 3’ UTR region. (**c**) Growth phenotype of Col-0, the *vps28-2* and *vps37-1.1* single and double (*vps28-2*/*vps37-1.1*) mutants, and *VPS28-2* and *VPS37-1*-overexpressing transgenic line plants. Photo was taken when the plants were approximately 5 weeks old.

**Supplementary Figure 16.**
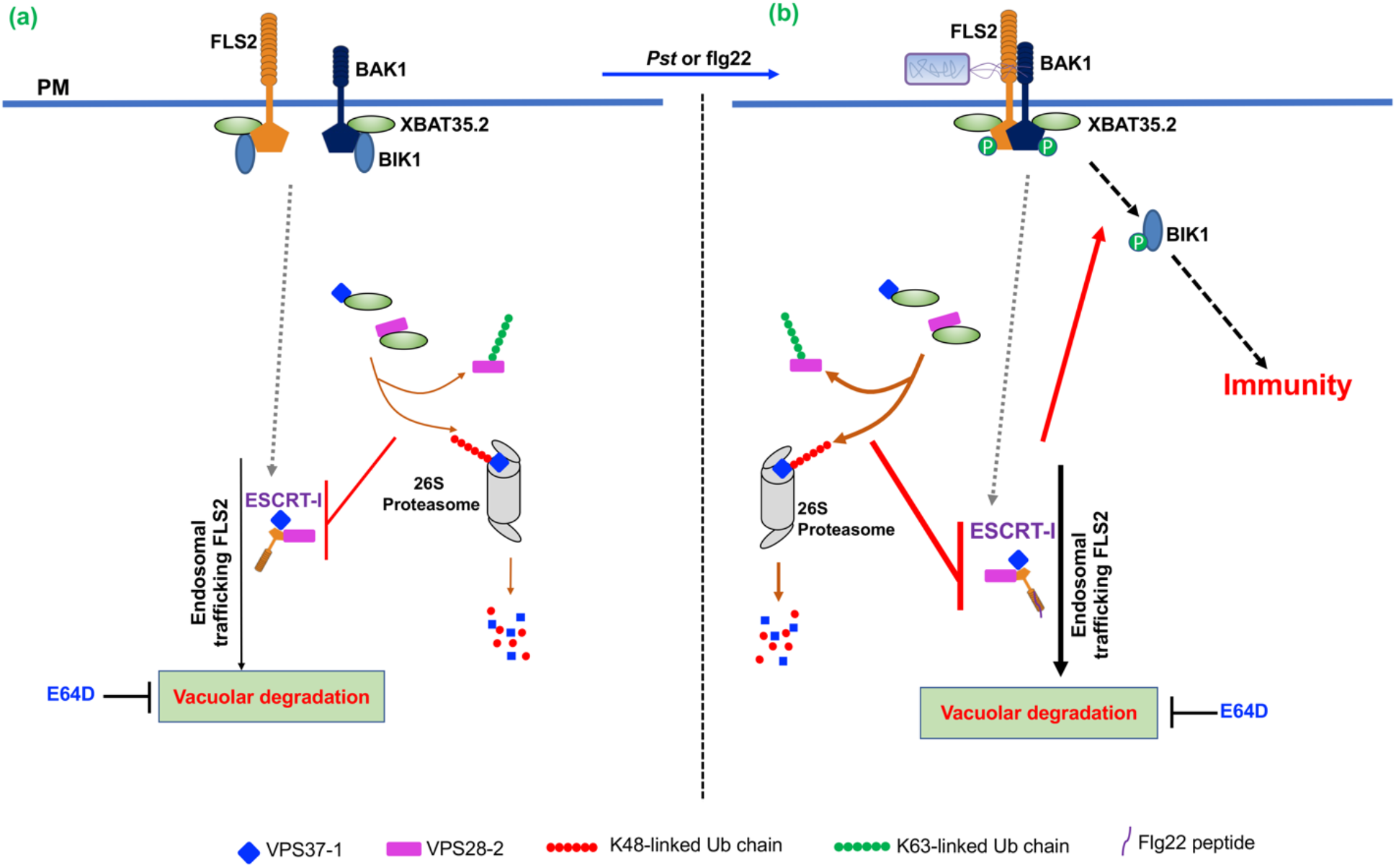
A working model depicting the positive regulation of FLS2-mediated plant immunity by XBAT35.2 through modifying the ESCRT-I subunits VPS28-2 and VPS37-1 with distinct linkage types of ubiquitin chains. **(a)** When the *Arabidopsis* plant is not infected by *Pseudomonas syringae* pv *tomato* (*Pst*) or treated with flg22, FLS2-mediated immunity is not activated. Under such circumstances, the Ankyrin repeats (ANK)-RING type ubiquitin E3 ligase XBAT35.2 is constitutively associated with FLS2, BAK1, and BIK1 and nonactivated FLS2 is constitutively endocytosed and recycled back to the PM (Beck et al., 2012). Our results suggest that a part of the endocytosed nonactivated FLS2 proteins are also targeted for ECSRT-I (VPS28-2 and VPS37-1)-mediated vacuolar degradation. **(b)** Upon *Pst* infection or flg22 treatment, FLS2 binds to bacterial flagellin or flg22, leading to the activation of FLS2 and immune signaling that culminates in plant immunity. Flg22 treatment enhances the association of XBAT35.2 with FLS2 and BAK1 but promotes the dissociation between XBAT35.2 and BIK1. The activated FLS2 bound with flg22 undergoes endocytosis and are targeted for vacuolar breakdown (Choi et al., 2013). The degradation of FLS2 at the vacuole is inhibited by E64D, an exogenous inhibitor of lysosomal/vacuolar hydrolases. XBAT35.2 sabotages VPS28-2/VPS37-1-mediated vacuolar degradation of both non-activate (**a**) and activated (**b**) FLS2 by catalyzing attachment of K48 and K63-linked polyubiquitin chains to VPS37-1 and VPS28-2, respectively, which leads to proteasomal degradation of VPS37-1 and disassociation of VPS28-2 from FLS2. The effect of stabilizing activated FLS2 by XBAT35.2 upon activation of plant immunity is more significant than that on non-activated FLS2 (as illustrated by thicker lines in **b**). By doing so, XBAT35.2 effectively stabilizes FLS2 and promotes host plant immunity against bacterial pathogens. The dotted arrows in gray indicate endocytosis and endosomal trafficking steps reported previously and not investigated in the present study.

**Supplementary Table 1.**
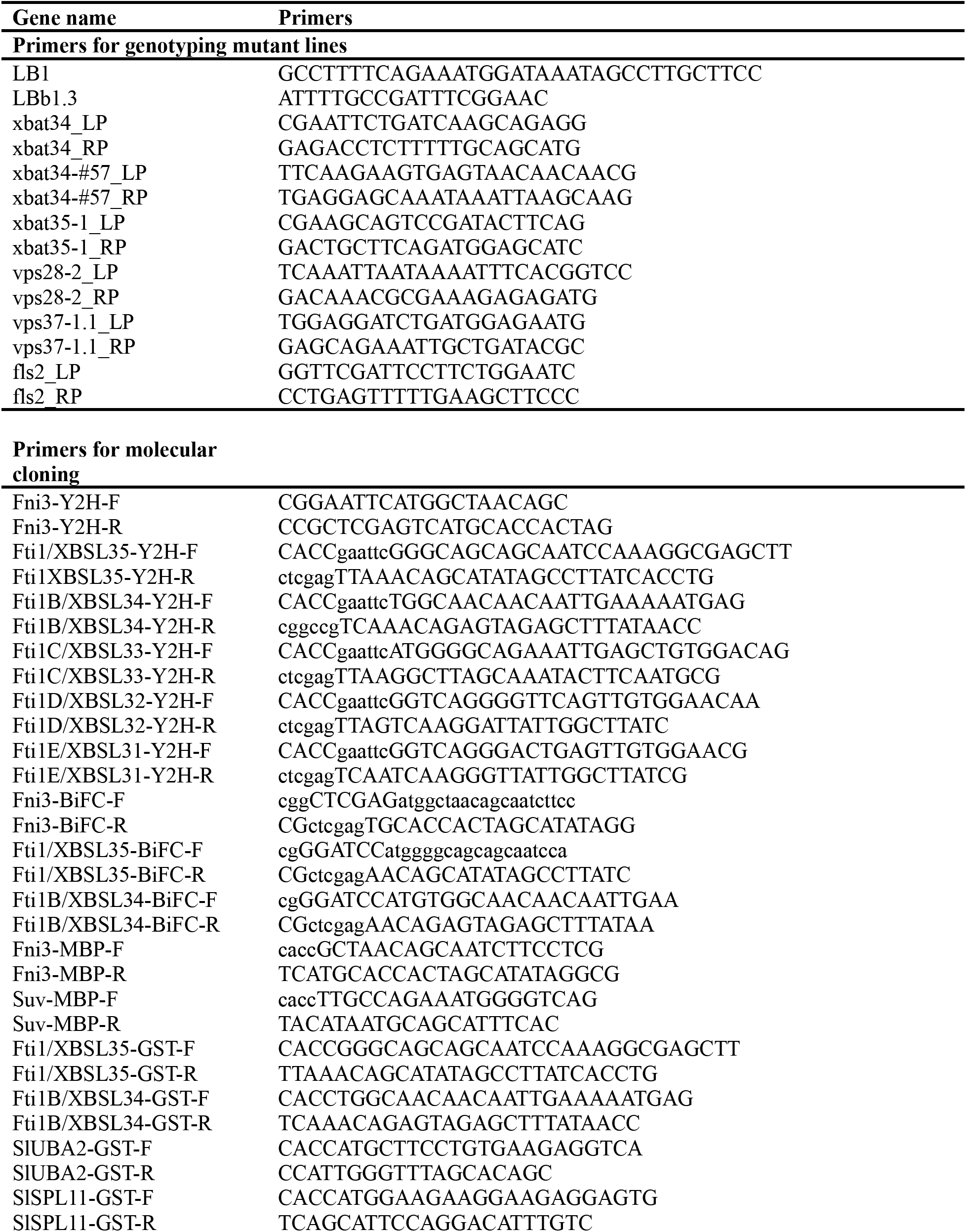
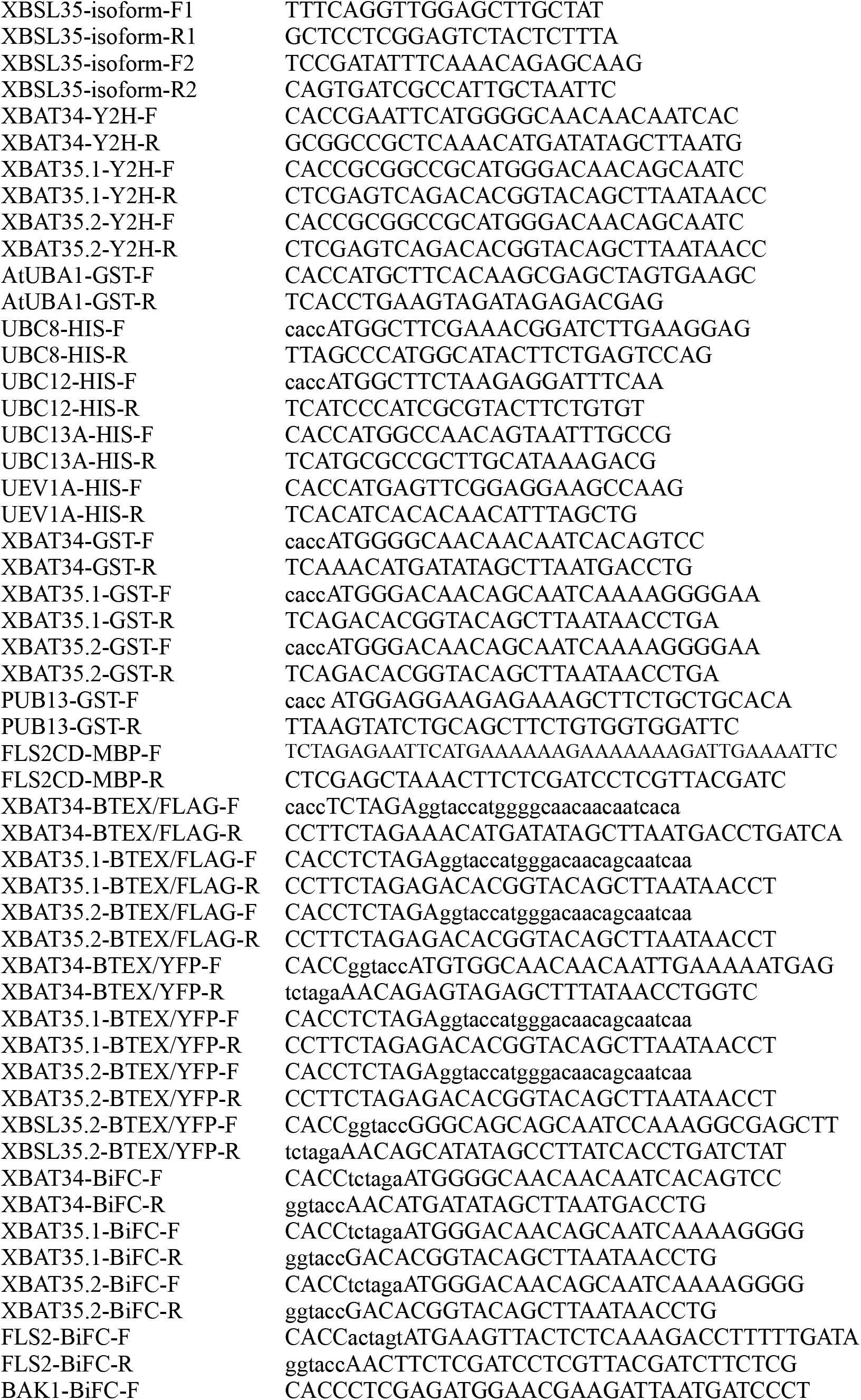
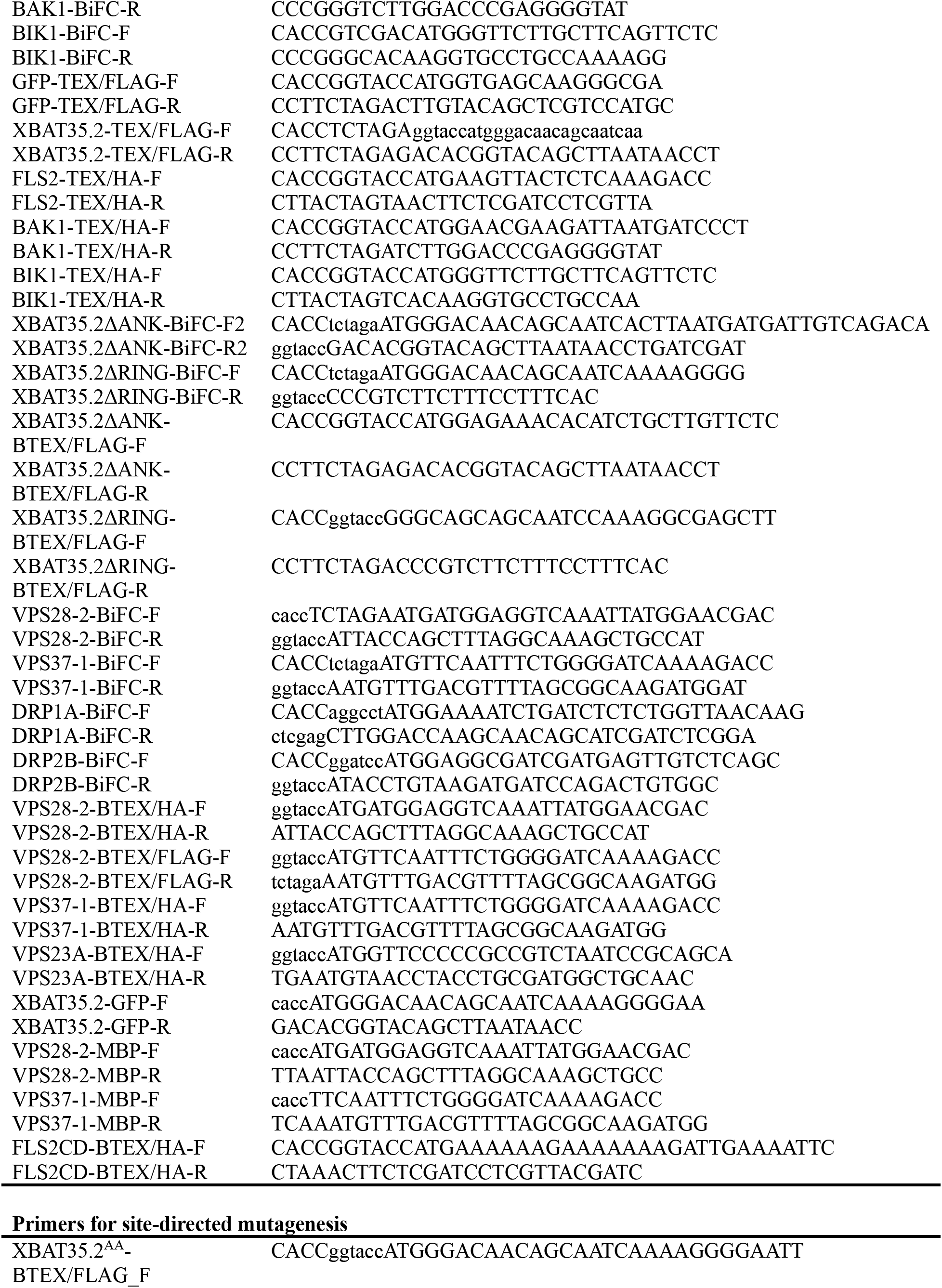
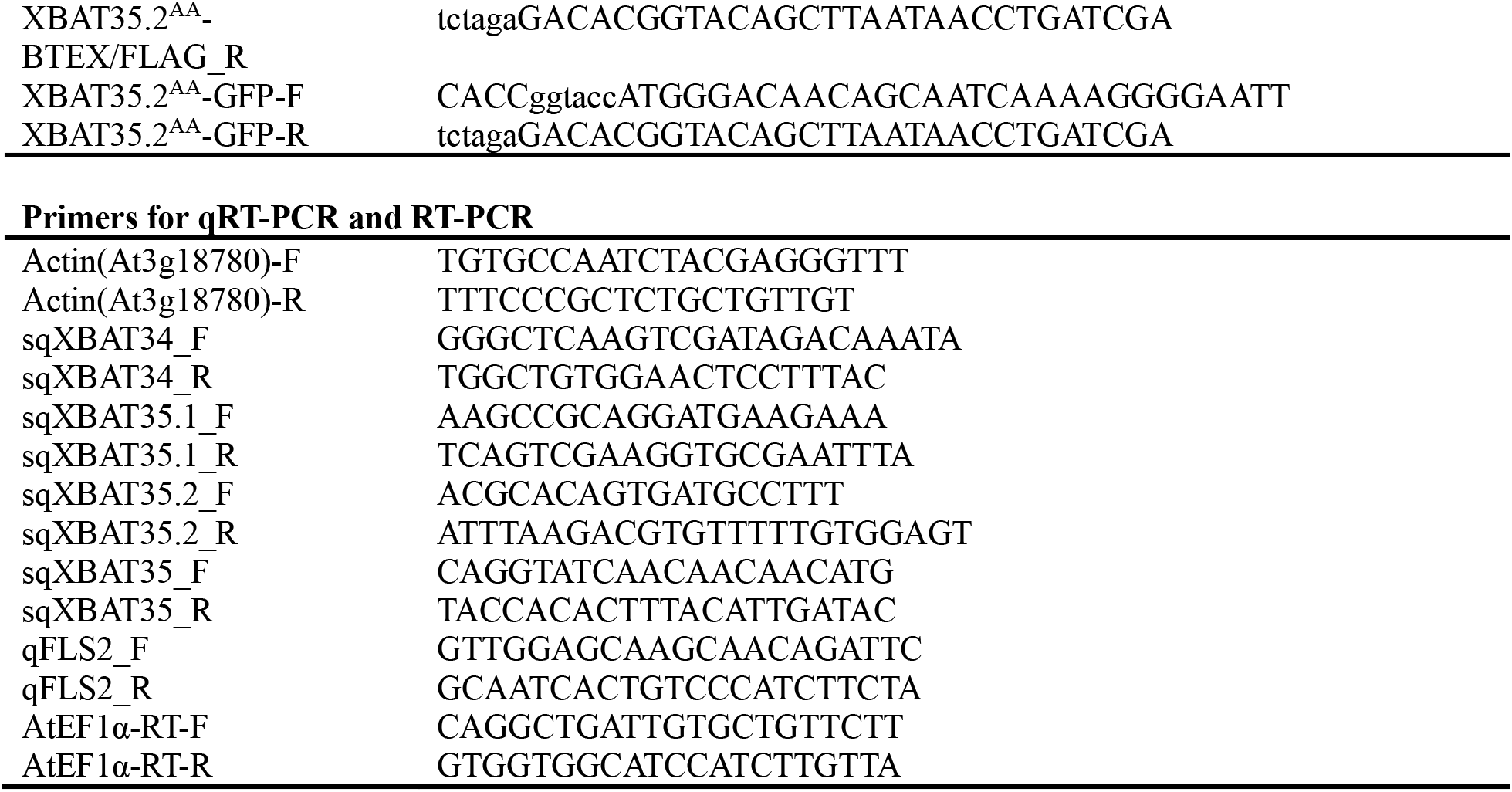
Primers used for genotyping, cloning, qRT-PCR, and other experiments.

